# Salt transport by the gill Na^+^-K^+^-2Cl^-^ symporter in palaemonid shrimps: exploring physiological, molecular and evolutionary landscapes

**DOI:** 10.1101/2020.04.30.070672

**Authors:** Anieli Cristina Maraschi, Samuel Coelho Faria, John Campbell McNamara

## Abstract

Palaemonid shrimps include species from distinct osmotic niches that hyper-regulate hemolymph osmolality and ionic concentrations in dilute media but hypo-regulate in saline media. Their gill epithelia express ion transporters like the Na^+^-K^+^-2Cl^-^ symporter (NKCC) thought to play a role in salt secretion. Using a palaemonid series from niches including marine tide pools through estuaries (*Palaemon*) to coastal and continental fresh waters (*Macrobrachium*), we established their critical upper salinity limits (UL_50_) and examined their short-(24 h) and long-term (120 h) hypo-regulatory abilities at salinities corresponding to 80% of the UL_50_’s (80%UL_50_). We tested for phylogenetic correlations between gill NKCC gene and protein expression and hemolymph Cl^-^ hypo-regulatory capability, and evaluated whether niche salinity might have driven gill NKCC expression. The *Palaemon* species from saline habitats showed the highest UL_50_’s and greatest hypo-regulatory capabilities compared to the *Macrobrachium* species among which UL_50_’s were higher in the diadromous than in the hololimnetic species. While basal gill NKCC mRNA transcription rates differed among species, expressions were unaffected by exposure time or salinity, suggesting post-transcriptional regulation of protein synthesis. Unexpectedly, hemolymph Cl^-^ *hyper*-regulatory capability correlated with gill NKCC gene expression, while gill NKCC protein synthesis was associated with *hyper*-regulation of hemolymph osmolality at the 80%UL_50_’s of almost all *Macrobrachium* species, suggesting a role for the gill NKCC symporter in salt uptake. The evolutionary history of osmoregulation in these palaemonid shrimps suggests that, while some molecular and systemic mechanisms have accompanied cladogenetic events during radiation into different osmotic niches, others may be driven by salinity.

## INTRODUCTION

The subphylum Crustacea includes innumerous species that have radiated into many different environments from their ancestral marine setting. The conquest of continental aquatic or terrestrial habitats has been enabled by adaptation to both abiotic and biotic factors, the subjacent physiological processes being mainly environmentally driven but also phylogenetically constrained (McNamara and Faria 2012, Faria et al. 2017, Faria et al. 2020).

For many crustaceans, the regulation of water and ion movements between the internal *milieu* and the external environment can constitute an energetically demanding challenge since their respective osmotic and ionic concentrations determine the intensity and direction of water and ion fluxes between the intra- and extracellular spaces, and between these and the external medium (Péqueux 1995, Freire et al. 2008a, McNamara and Faria 2012). Osmotic and ionic homeostasis thus constitutes one of the essential physiological determinants of niche diversification since the evolution of osmoregulatory traits can dictate the degree of euryhalinity of a species and thus affects the potential for habitat diversification (Lee et al. 2011, Faria et al. 2011, McNamara and Faria 2012, Velotta et al. 2016).

The caridean shrimp family Palaemonidae has diversified into most aquatic niches, ranging from osmotically stable environments like the fully marine and freshwater biotopes to habitats exhibiting daily and seasonal variations in salinity such as estuarine and intertidal biotopes (Freire et al. 2003, Augusto et al. 2009). Many diadromous species also make ontogenetic transitions as zoeae and post-larvae among fresh, brackish and marine waters (Moreira et al. 1983, Freire et al. 2003, Boudour-Boucheker et al. 2014). The occupation of environments of differing salinity regimes can generate specific adaptations at distinct levels of phylogenetic grouping, such as systemic abilities to regulate hemolymph osmolalities and ion concentrations (Freire et al. 2008a, McNamara and Faria 2012, Thabet et al. 2017).

Species from dilute and freshwater habitats, particularly representatives of the genera *Macrobrachium* and *Palaemonetes*, show reduced gill epithelial permeability to water and ions, accompanied by active salt absorption predominantly by the gills (Péqueux 1995, Kirschner 2004, Freire et al. 2008a, McNamara and Faria 2012), and water excretion as urine by the antennal glands (McNamara et al. 2015). The ability to hypo-regulate hemolymph osmolality seen in marine, estuarine or brackish water species of *Palaemon* is characterized by active epithelial salt secretion by the gills, in contrast to hyper-osmoregulation effected when in dilute media. Such hyper-/hypo-osmoregulatory activity furnishes a fairly constant osmotic and ionic concentration and volume of the extracellular fluid, independently of environmental salinity, a process known as anisosmotic extracellular regulation (Péqueux 1995, Freire et al. 2008a, McNamara and Faria 2012, Thabet et al. 2017).

The species of the genus *Macrobrachium* are strong hyper-osmoregulators and maintain elevated osmotic and ionic gradients (≈30: 1) when in fresh water (Δ_hemolymph/external medium_ ≈400 mOsm/kg H_2_O; Moreira et al. 1983, Faleiros et al. 2010, Faria et al. 2011, Maraschi et al. 2015, Freire et al. 2018), owing to gill salt uptake associated with reduced epithelial permeability (Freire et al. 2008a, McNamara and Faria 2012, McNamara et al. 2015). Reconstruction of the ancestral palaemonid environment suggests the origin of a weakly hyperosmotic regulator (660 mOsm/kg H_2_O) that inhabited an estuarine or brackish water niche (510 mOsm/kg H_2_O, 17 ‰ salinity) (McNamara and Faria 2012). Further, the reduced hemolymph osmolalities characteristic of the *Macrobrachium* lineage (McNamara and Faria 2012) appear to constitute an adaptive condition subsequently inherited by the more derived groups. Some species of freshwater *Macrobrachium* exhibit notable hypo-osmoregulatory ability in addition to their hyper-osmoregulatory capacity (Moreira et al. 1983, Freire et al. 2003), although limited compared to the considerable hypo-regulatory capability of species from variable salinity habitats like *Palaemon northropi* and *P. pandaliformis* (Freire et al. 2003, Faleiros et al. 2017). This is likely the result of the loss of the adaptive value of salt secretion in freshwater species (McNamara and Faria 2012, McNamara et al. 2015). While osmoregulatory mechanisms have been widely explored in different crustacean groups, the mechanisms of salt secretion and their evolution among the palaemonid shrimps are entirely unknown (see McNamara and Faria 2012).

The different patterns of osmoregulatory ability seen among the Crustacea derive mainly from distinct arrangements of gill epithelial ion transporters that vary from marine osmoconformers to brackish and freshwater hyper-regulators (McNamara and Faria 2012). In these latter species, gill salt uptake requires the combined action of electrogenic ion pumps like the V(H^+^)-ATPase (VAT) and the Na^+^/K^+^-ATPase (NKA) to generate electrochemical gradients and the consequent absorption of Na^+^ ions across apical Na^+^ channels and the Na^+^/H^+^ exchanger, and Cl^-^ ions through the apical Cl^-^/HCO_3_^-^ anti-porter. The H^+^ and HCO_3_^-^ ions are generated by the reversible hydration of metabolic CO_2_ by intracellular carbonic anhydrase (Péqueux 1995, Kirschner 2004, Freire et al. 2008a, McNamara and Faria 2012, Maraschi et al. 2015).

The mechanism of salt secretion responsible for hypo-osmoregulation in palaemonid shrimps from marine or hyperosmotic environments, and of chloride hypo-regulation in freshwater species, is far from clear. Based on vertebrate models (Hwang 2009, Evans et al. 2010, Gonzalez 2012, Hiroi and McCormick 2012), transcellular Cl^-^ transport by the sodium-potassium-two chloride symporter (NKCC) and chloride channels, driven partly by the basal NKA, together with paracellular Na^+^ efflux, is thought to mediate salt secretion (McNamara and Faria 2012). The NKA, located in the membrane invaginations of the intralamellar ionocytes (McNamara and Torres 1999), transports three Na^+^ ions to the hemolymph in exchange for two K^+^ ions to the intracellular fluid (Leone et al. 2017). The resulting inward Na^+^ gradient generated by the NKA together with passive Na^+^ influx to the hemolymph from the external medium, drives the flow of Na^+^, and particularly of Cl^-^, through the NKCC symporter, putatively inserted in the ionocyte invaginations or in the lower pillar cell flange cell membranes. Chloride efflux occurs through apical channels in the pillar cells, generating a negative external transepithelial voltage, which drives the paracellular efflux of Na^+^ (Freire et al. 2008a McNamara and Faria 2012). K^+^ is recycled to the hemolymph via basal K^+^ channels, and Na^+^ through the NKA.

Although still rare, some molecular findings have correlated expression of the NKCC symporter gene with salt secretion and/or absorption. Two major NKCC isoforms have been identified in vertebrates: NKCC1 and NKCC2, with distinct functions in salt secretion and reabsorption, respectively (Hass and Forbush 2000). With regard to crustacean osmoregulation, molecular studies on membrane transporter localization, activity and expression are helping to elucidate the mechanisms of salt transport (Péqueux 1995, Kirschner 2004, Freire et al. 2008a, McNamara and Faria 2012). NKCC mRNA expression increases in the posterior gills of the semi-terrestrial estuarine crab *Neohelice granulata* exposed to either dilute or concentrated seawater (Luquet et al. 2005) in contrast to the swimming crab *Portunus trituberculus* in which expression is down regulated (Lv et al. 2016). NKCC mRNA expression is up regulated in the gills of the hololimnetic freshwater shrimp *Macrobrachium australiense* when challenged by increased salinity (Moshtaghi et al. 2018). Gill NKCC gene expression has been characterized in different molt cycle phases in the estuarine mangrove crab *Scylla paramamosain*, revealing an important role for the symporter in salt uptake during the post-molt stages (Xu et al. 2017).

Here, we investigate hypo-osmotic and particularly Cl^-^ hypo-regulation in different species of *Palaemon* and *Macrobrachium*, exhibiting different life histories, habitats and osmotic niches. Couched within a phylogenetic framework, we accompany gene and protein expression of the gill sodium-potassium-two chloride symporter, examining their relationship with Cl^-^ hypo-regulatory capability, and appraising whether niche salinity might have constituted a driver of NKCC expression during palaemonid shrimp radiation. This approach allows an objective evaluation of the roles of salinity-driven traits and shared ancestry in the physiological evolution of osmoregulatory processes.

## MATERIAL AND METHODS

Adult, non-ovigerous, intermolt, male and female palaemonid shrimps, measuring from 3 to 5 cm total length depending on the species investigated, were collected from continental streams and reservoirs, coastal estuaries and river mouths, and from tide pools at low tide, in southeastern Brazil. The shrimps were captured manually by sieving the marginal vegetation using a builder’s sieve or leaving baited plastic traps overnight. Collections were authorized under ICMBio/MMA permit #29594-12.

*Palaemon pandaliformis* (Stimpson 1871), an estuarine shrimp, was collected from an estuary in Paraná State (25° 34’ 23.64” S, 48° 21’ 3.43” W). *Macrobrachium potiuna* (Müller 1880), a hololimnetic freshwater shrimp, was collected from a continental stream also in Paraná State (25° 31’ 01.25” S, 49° 00’ 30.55” W). Shrimps were transported in aerated 25-L carboys containing water from their respective collection sites to the Laboratory of Comparative Osmoregulatory Physiology, Federal University of Paraná, Curitiba, Paraná State. These two species were acclimatized to laboratory conditions in 60-L plastic tanks containing dilute seawater (17 ‰S, g/L, salinity) or fresh water (<0.5 ‰S), respectively, under constant aeration for 5 days at room temperature.

*Palaemon northropi* (Rankin 1898), a tide pool shrimp, was collected from rocky coastal shores in São Paulo State (23° 49’ 53.44” S, 45° 31’ 16.58” W). *Macrobrachium acanthurus* (Wiegmann 1836) and *Macrobrachium olfersii* (Wiegmann 1836), diadromous freshwater shrimps, were collected from the marginal vegetation of nearby coastal streams (Guaecá river, 23° 49’ 26.22” S; 45° 27’ 8.66” W and the Paúba river, 23° 47’ 51.90” S; 45° 32’ 39.59” W, respectively) while *Macrobrachium amazonicum* (Heller 1862), also diadromous, was collected from a land-locked population found in an inland reservoir in northeastern São Paulo State (21° 06’ 34.78” S; 48° 03’ 06.51” W). *Macrobrachium brasiliense* (Heller 1862), a hololimnetic freshwater shrimp, was collected from a continental stream (21° 20’ 17.35” S; 47° 31’ 22.08” W) in the same region.

All species were transported in aerated 25-L carboys containing water from their respective collection sites to the Laboratory of Crustacean Physiology, University of São Paulo, Ribeirão Preto, São Paulo State. The *Macrobrachium* species were acclimatized to laboratory conditions in 60-L plastic tanks containing fresh water from the collection sites (<0.5 ‰S) while *P. northropi* was held in dilute seawater (18 ‰S), under constant aeration for 5 days at room temperature. All shrimps were fed every 48 h with fish fragments and grated carrot. Holding tanks were cleaned weekly when the water was changed.

### Survival, upper critical salinity limits and experimental protocol

#### Survival

A parameter commonly used to characterize salinity tolerance is the ‘upper critical salinity limit’ (UL_50_), which corresponds to the lethal salt concentration that induces 50% mortality during an arbitrary exposure period (Thurman 2002, 2003, Kefford et al. 2004, Faria et al. 2017). Using the UL_50_ for each species ensures that the relative degree of experimental osmotic challenge is comparable among the different species.

We opted to challenge each species with a carefully delimited, ecologically relevant, salinity range rather than to employ a ‘common garden’ experimental design using the same range for all species, given the different osmotic niches occupied naturally by each.

To establish their respective UL_50_’s, eight specimens of each palaemonid shrimp species were separated into groups of two individuals each in four 2-L aquaria for each experimental salinity, and were exposed for 5 days (*i. e*., a total of N=8 shrimps per species, with four replicates). Salinity ranges were species specific and varied from 16 to 50 ‰S. All aquaria were provided with constant aeration and held at room temperature (23 °C). Survival was checked every 12 h and shrimps that did not recover their initial position after being placed in lateral *decubitus* were considered ‘dead’.

UL_50_’s for each species were estimated by Probit analyses that adjusted the respective percentage survival rates to a linear regression (Finney 1971, Thurman 2002, 2003). The UL_50_ salinity values were 43.1 ‰S for *P. northropi*, 39.7 for *P. pandaliformis*, 31.4 for *M. acanthurus*, 28 for *M. olfersii*, 24.7 for *M. brasiliense*, 24.5 for *M. amazonicum* and 24.0 ‰S for *M. potiuna*. To avoid excessive mortality, the hyperosmotic salinity challenges employed here correspond to 80% of the estimated UL_50_ values (80%UL_50_), *i. e*., 35 ‰S for *P. northropi*, 32 for *P. pandaliformis*, 25 for *M. acanthurus*, 22 for *M. olfersii*, 20 for *M. amazonicum* and *M. brasiliense*, and 19 ‰S for *M. potiuna*.

The time courses of salinity challenge at these 80%UL_50_’s were performed as described above for establishing the UL_50_’s, using 24-h or 120-h exposure periods.

### Measurement of hemolymph osmolality and chloride concentration

Specimens (N = 8) of each shrimp species were exposed for 24 or 120 h to the respective 80%UL_50_ salinity from the initial laboratory acclimatization salinity (Time = 0 h). After briefly chilling in crushed ice, a hemolymph sample was drawn from the pericardial sinus of each shrimp (N = 8) through the arthrodial membrane between the posterior margin of the cephalothorax and the first abdominal segment using a #27-5 needle coupled to an insulin syringe.

Hemolymph osmolality was measured in 10-μL aliquots using a vapor pressure micro-osmometer (Wescor Vapro 5600, Logan UT, USA). Hemolymph chloride was titrated in 10-μL aliquots against mercury nitrate using s-diphenylcarbazone as the indicator (Schales and Schales 1941, modified by Santos and McNamara 1996) with a microtitrator (Model E485, Metrohm AG, Herisau, Switzerland).

### Muscle water content

Abdominal muscle fragments of ≈50-100 mg fresh mass each were dissected and the excess fluid was blotted off. Samples were placed in tared micro-Eppendorf tubes and weighed immediately on a precision analytical balance (Ohaus AP250D, Parsippany, New Jersey, USA, ±10 µg precision). The tubes containing the fragments were then oven dried at 60 °C for 24 h to obtain the sample dry masses. Muscle water content (%) was calculated as [(fresh mass - dry mass)/fresh mass] × 100.

### RNA extraction and amplification of the gill ribosomal protein L10 and sodium-potassium-two chloride symporter partial cDNA sequences

All 7 gills from each individual shrimp (5 ≤ N ≤ 8) were dissected under a magnifying lens with micro-scissors and homogenized in TRIzol reagent (Life Technologies, Thermo Fisher Scientific, Waltham, MA, USA) (1: 100 w/v) following the manufacturer’s protocol. The purity of the total RNA in the gill samples, extracted under RNAse free conditions, was evaluated from the absorbance ratios at 260 and 280 nm (Qubit 2.0 fluorometer, Thermo Fisher Scientific). Homogenate samples were stored at −80 °C until use, either for sequencing or for quantitative gene expression.

Gill sample RNA was treated with DNAse I (Life Technologies) to ensure the absence of contaminating DNA, and was evaluated by PCR, using primers for the ribosomal protein L10 gene (RPL10Cs_F and RPL10Cs_R, see Table 1), followed by confirmation with 1% agarose gel electrophoresis. The gill RNA samples were then submitted to a cDNA synthesis reaction by PCR (Veriti Thermal Cycler, Life Technologies) using the SuperScript III RT-PCR kit (Life Technologies), employing reverse transcriptase in the presence of deoxynucleotides (dNTP, Life Technologies) in RT-III buffer solution. The thermocycling protocol was: 5 min at 94 °C, followed by 40 cycles of 5 s each at 94 °C, 45 s at 40 °C (for RPL10) or 55 °C (for NKCC from *M. amazonicum*) or 50 °C (for NKCC from all other palaemonids), 1 min at 72 °C, and a final cycle for 10 min at 72 °C. All samples were verified in 1% agarose gels for successful cDNA amplification by PCR using the RPL10Cs_F and RPL10Cs_R primers.

**Table 1.**
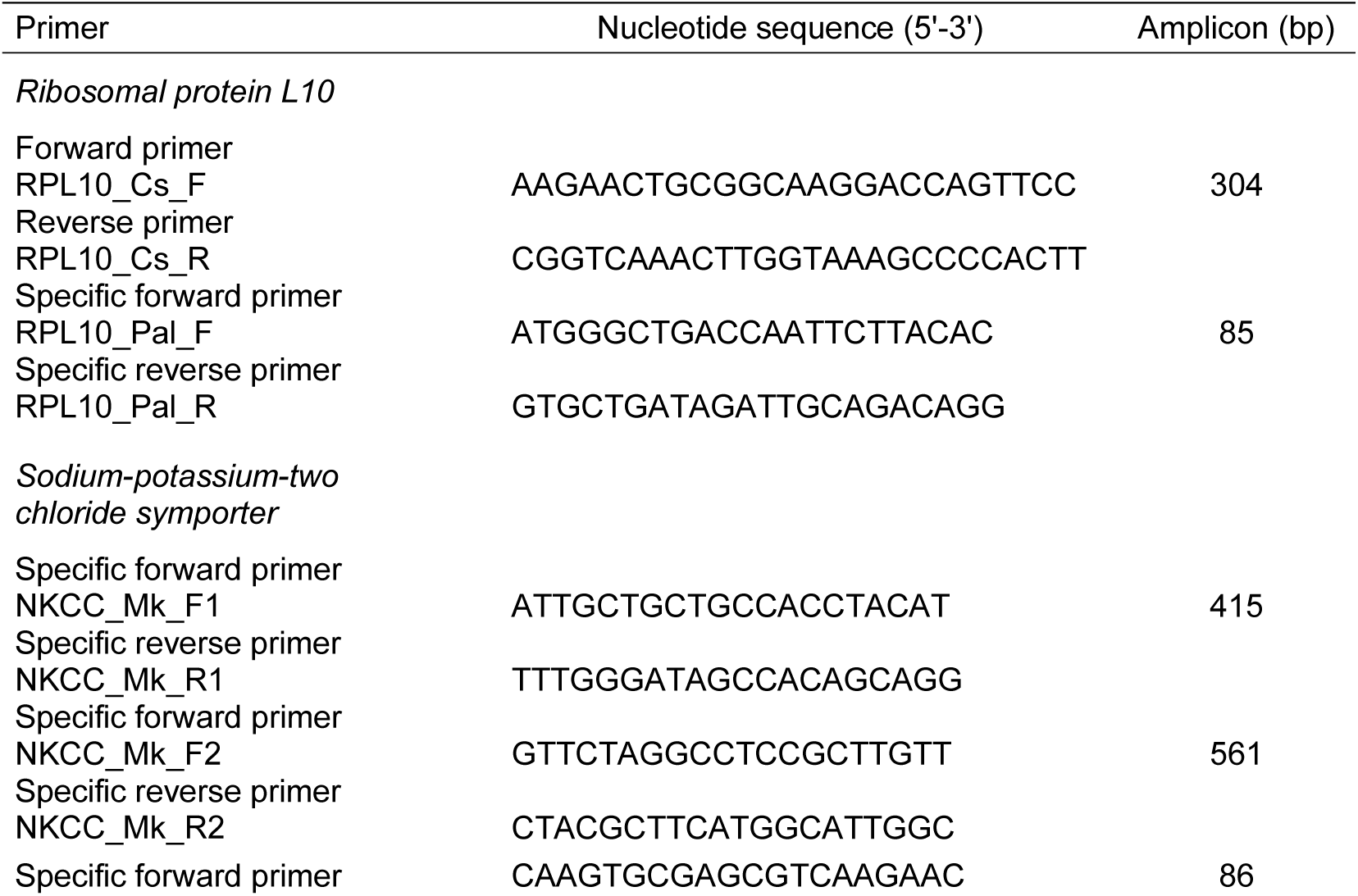
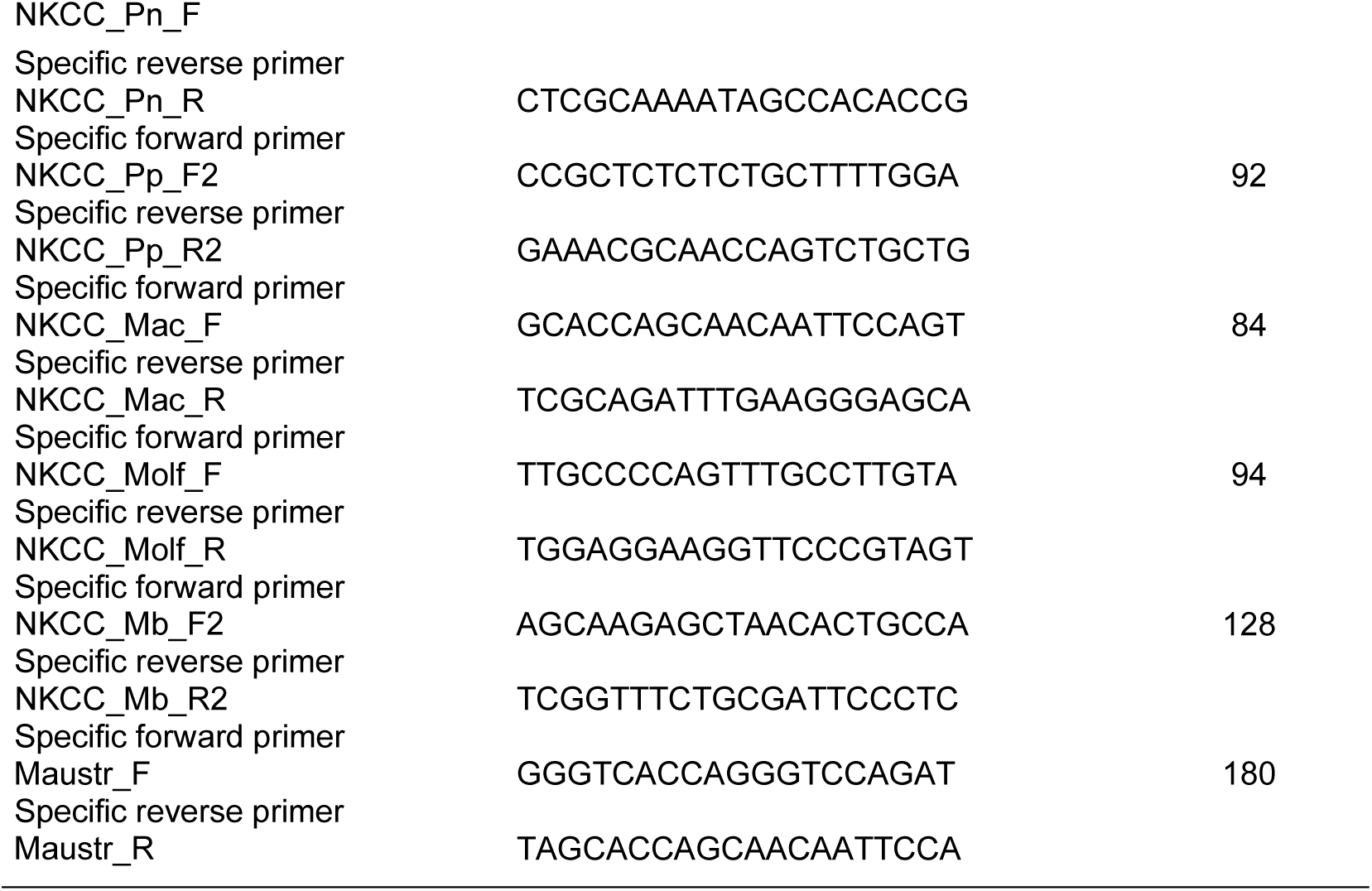
Characteristics of the forward and reverse primer pairs used to partially amplify the ribosomal protein L10 (RPL10) and sodium-potassium-two chloride symporter (NKCC) genes, and to quantify their expression in palaemonid shrimp gills. Primers for the RPL10 reference gene were based on the *Callinectes sapidus* cDNA sequence (RPL10_Cs_F/R, Wynn et al., 2004) and on a multiple alignment of the *Macrobrachium acanthurus, M. amazonicum* and *M. brasiliense* cDNA sequences (RPL10_Pal_F/R, Faleiros et al., 2010). Primer pairs used to partially amplify the NKCC gene in *M. acanthurus, M. amazonicum, M. brasiliense* and *M. potiuna* (NKCC_Mk_F1/R1) and in *M. olfersii, Palaemon northropi* and *P. pandaliformis* (NKCC_Mk_F2/R2) were based on the complete gill NKCC sequence for *M. koombooloomba*, kindly provided by Dr David A Hurwood (QUT, Australia). Specific primer pairs used to quantify gill NKCC gene expression were based on the partial cDNA sequences of the NKCC gene obtained in *P. northropi* (NKCC_Pn_F/R), *P. pandaliformis* (NKCC_Pp_F2/R2), *M. acanthurus* and *M. amazonicum* (NKCC_Mac_F/R), *M. olfersii* (NKCC_Molf_F/R), *M. brasiliense* (NKCC_Mb_F/R2) and *M. potiuna* (NKCC_Maustr_F/R, based on *M. australiense*, Moshtaghi et al., 2018).

The primers used to partially amplify the gill RPL10 gene sequence (RPL10_Cs_F and RPL10_Cs_R) in the seven palaemonid species were designed based on a conserved sequence from the *Callinectes sapidus* gene, obtained from GenBank (AY822650, Wynn et al. 2004) (see Table 1). The RPL10 gene encodes for ribosomal protein L10 and was used as an endogenous reference control for the quantitative gene expression experiments.

The two primer pairs used for partial amplification of the gill NKCC gene in the seven target species were based on the complete sequence for this gene in *Macrobrachium koombooloomba* (kindly provided by Dr David A Hurwood, Queensland University of Technology, Australia) and on a sequence for the gill NKCC from *M. australiense* (Rahi et al. 2017) (see Table 1).

### Cloning and sequencing of partial gill ribosomal protein L10 and sodium-potassium-two chloride symporter genes

The fragments obtained from selected PCR amplifications were cut from the gel, extracted and purified with the PureLink Quick Gel Extract Kit (Life Technologies), cloned into the PCR TOPO TA vector (Life Technologies) and transformed in a thermo-competent *Escherichia coli* DH5α bacterial strain.

Plasmid DNA samples from several selected successful clones containing the RPL10 and NKCC inserts were purified with the PureLink Plasmid Mini Kit (Life Technologies) and sequenced using the traditional method of incorporating dideoxynucleotides (Sanger et al. 1977), employing the universal primers M13F and M13R. After sequencing the amplified cDNA clones, the identity of the nucleotide sequences and of the predicted translated amino acid sequences of the corresponding proteins were analyzed using the NCBI BLAST algorithm (Altschul et al. 1990; https://blast.ncbi.nlm.nih.gov) which detects regions of similarity between sequences and compares them with known sequences deposited with public databases. All partial nucleotide sequences generated here for the RPL10 and NKCC genes were deposited with NCBI GenBank (https://www.ncbi.nlm.nih.gov/genbank/).

### Quantitative expression of the ribosomal protein L10 and sodium-potassium-two chloride symporter genes

Quantitative PCR reactions were performed in triplicate for each of the 5-8 gill samples obtained for each species at each of the three time intervals (0, 24 and 120 h) at their respective 80%UL_50_’s using the PowerUp SYBR Green PCR Master Mix Kit (LifeTechnologies), following the manufacturer’s instructions, employing a BioRad CFX96 C1000 Touch Real-Time PCR Detection System (Hercules, CA, USA).

The thermocycling protocol was: an initial step of 10 min at 95 °C, 40 cycles of 15 s each at 95 °C, and a 1-min cycle at 60 °C. After the reaction was completed, a dissociation or melting curve was performed to check for contaminants, formation of primer dimers and/or amplification of more than one amplicon. The thermocycling protocol for constructing the dissociation curve was: an initial cycle of 5 s at 65 °C, 60 cycles of 5 s each increasing by 0.5 °C every cycle up to 95 °C, 5 s at 95 °C.

The comparative CT method was used to quantify gill NKCC gene expression in the samples from the different species at the different exposure times. A standard curve validation was performed to evaluate the similarity between the amplification efficiencies [E = 10^(−1/slope)^] of the target gill NKCC genes and the endogenous RPL10 control genes. Our adjusted curve (R^2^ > 0.996) corresponded to efficiencies between 90 and 110%. The comparative CT method uses the exponential formula (2^-ΔΔCT^) to quantify target gene expression in the samples (Livak and Schmittgen 2001), which are then normalized by expression of the constitutive RPL10 gene in the same sample.

A standard curve for each primer pair (see Table 1) for each species was constructed from the serial dilution of a pool of cDNAs. The dilution used for individual cDNA samples was determined from the values that best fit the curve. Negative controls (No Template Control) were performed without the cDNA template to detect possible contamination.

### Quantification of gill sodium-potassium-two chloride protein expression by Western blotting

Total protein was extracted from the gill homogenates of all species under all experimental conditions during processing with TRIzol reagent to obtain total RNA, according to the manufacturer’s instructions. The protein phases were isolated, resuspended in 1% SDS and stored at −20 °C until assay. Quantification of total protein was evaluated using the Pierce™ BCA Protein Assay kit (Thermo Fisher Scientific), employing an ELISA microplate reader (SpectraMax M2, Molecular Devices LLC, San Jose, CA, USA) at 562 nm absorbance.

Aliquots of the protein extract containing 20 μg total protein were diluted in 10 µL of sample buffer (2: 1 v/v) and were submitted to gel electrophoresis in10% acrylamide in 150 mM Tris-Base buffer containing 1 mM SDS (pH 8.8). The stacking gel was 5% acrylamide in 50 mM Tris-Base buffer containing 1 mM SDS (pH 6.8). Electrophoresis was performed in Tris-glycine buffer (pH 8.3) containing 0.1% SDS during ≈4 h at 20 mA. The proteins and peptides in the gel were transferred overnight at 4 °C and 135 mA to a Hybond-P PVDF membrane (Amersham Biosciences, Global Life Sciences Solutions LLC, Marlborough, MA, USA) previously incubated in 100% methanol followed by UltraPure water (Invitrogen, Thermo Fisher Scientific) in a Western blotting chamber containing a transfer buffer consisting of 25 mM Tris-Base, 192 mM glycine, 10% SDS and 20% methanol.

The membrane was incubated in an oven at 37 °C for 1 h and was rehydrated in 100% methanol, followed by UltraPure water and Tris-buffered saline with Tween (TBST). The membrane was then incubated in TBST containing 5% non-fat milk powder for 1 h to block non-specific binding sites, and cut in half at the mid line, corresponding to the point at which the 70-kDa proteins are separated. The upper half was incubated with a 131-kDa anti-NKCC primary antibody (mouse, clone T4 anti-IgG antibody, Developmental Studies Hybridoma Bank, University of Iowa, Iowa City IA, USA) and the lower half with a primary 42-kDa anti-β-actin antibody (chicken, ab-5, anti-mouse IgG, BD Biosciences, San Jose, CA, USA). Both antibodies were diluted 1: 1,000 in 0.1% bovine serum albumin (Sigma-Aldrich, St. Louis, MO, USA) + TBST and incubated for 2 h at room temperature.

The membrane was washed in TBST and incubated with a peroxidase-conjugated secondary antibody (goat, anti-mouse IgG, Sigma-Aldrich) diluted 1: 1,000 in TBST + 0.1% BSA for 2 h at room temperature, rewashed in TBST and covered with a DAB peroxidase chromogen solution (SigmaFAST 3,3’-Diaminobenzidine peroxidase substrate, Sigma-Aldrich) until the complete appearance of the protein bands of interest. After development, the membrane was immersed in UltraPure water to interrupt the reaction.

The membrane was photo-documented and protein abundance was quantified by analyzing the relative band intensity using ImageJ software (Schneider et al. 2012). Differential expression values of the sodium-potassium-two chloride symporter protein were normalized by comparison with the expression of β-actin in the same sample.

### Statistical analyses

#### Intra-specific analyses

After ascertaining the normality of the data distributions and the homogeneity of their variances, the effect of exposure time at each 80%UL_50_ salinity was evaluated on: (i) hemolymph osmolality and chloride concentration; (ii) gene expression of the gill sodium-potassium-two chloride symporter; and (iii) protein abundance of the gill sodium-potassium-two chloride symporter. The data were evaluated using one-way analyses of variance followed by the Student-Newman-Keuls (SNK) *post hoc* multiple means procedure to detect significant differences. A minimum significance level of P = 0.05 was employed for all tests. Analyses were performed using Sigma Plot 11.0 (Systat Software, Inc., San Jose, CA, USA), and the data are given as the mean ± standard error of the mean.

#### Phylogenetic comparative analyses

For all comparative analyses, we employed Pereira’s (1997) morphological phylogeny for palaemonid shrimps, assuming arbitrary branch lengths (Grafen 1989) and thus ensuring correct standardization of the phylogenetically independent contrasts (Felsenstein 1985).

The phylogenetic signal for each physiological trait was evaluated using Moran’s I autocorrelation analysis, which indicates the propensity for trait similarity between closely related species (Rezende and Diniz-Filho 2012). Moran’s I ranges from −1 to +1, significant positive values indicating similarity between closely related species, and significant negative values indicating dissimilarity (Gittleman et al. 1996). The analysis was performed using the Phylogenetic Analysis in Macroecology package (PAM version 0.9 beta, Rangel and Diniz-Filho 2012).

A phylogenetic, generalized least squares linear model (pGLS) was used to test the effect of 80%UL_50_ salinity on the physiological parameters and on the gene and protein expressions of the sodium-potassium-two chloride symporter in the shrimp gills. This model includes phylogenetic structure as a covariance matrix in the linear model, assuming phylogenetic dependence of the data (Grafen et al. 1989, Garland and Ives 2000, Lavin et al. 2008). Phylogenetic paired t-tests were used to evaluate differences between the different exposure times at the 80%UL_50_ salinity for the gene and protein expressions, thus including the phylogenetic non-independence of species (Lindenfors et al. 2010).

The comparative analyses were performed using the R Statistical Platform (R Development Core Team, 2009) with the *nlme* (Pinheiro et al. 2019) and *ape* packages (Paradis et al. 2004). A minimum significance level of P = 0.05 was used throughout.

## RESULTS

### Survival

The survival curves of the seven palaemonid species from habitats of different salinity during osmotic challenge up to 120 h are provided in Figure 1. The *Palaemon* species were more tolerant of increased salinity, mortality occurring at 45 ‰S in *P. northropi* and 35 ‰S in *P. pandaliformis*. Their respective upper salinity limits (UL_50_) were 43.1 ‰S and 39.7 ‰S. The diadromous *Macrobrachium* species were more salinity tolerant than the hololimnetic species. Mortality began at 30 ‰S in *M. acanthurus*, at 26 ‰S in *M. olfersii* and at 22 ‰S in *M. amazonicum*. UL_50_’s were 31.4 ‰S in *M. acanthurus*, 28 ‰S in *M. olfersi* and 24.5 ‰S in the land-locked *M. amazonicum*. Mortality in the hololimnetic species began at 22 ‰S in *M. potiuna* and 19 ‰S in *M. brasiliense*, with UL_50_’s of 24 ‰S and 24.7 ‰S, respectively.

**Figure 1.**
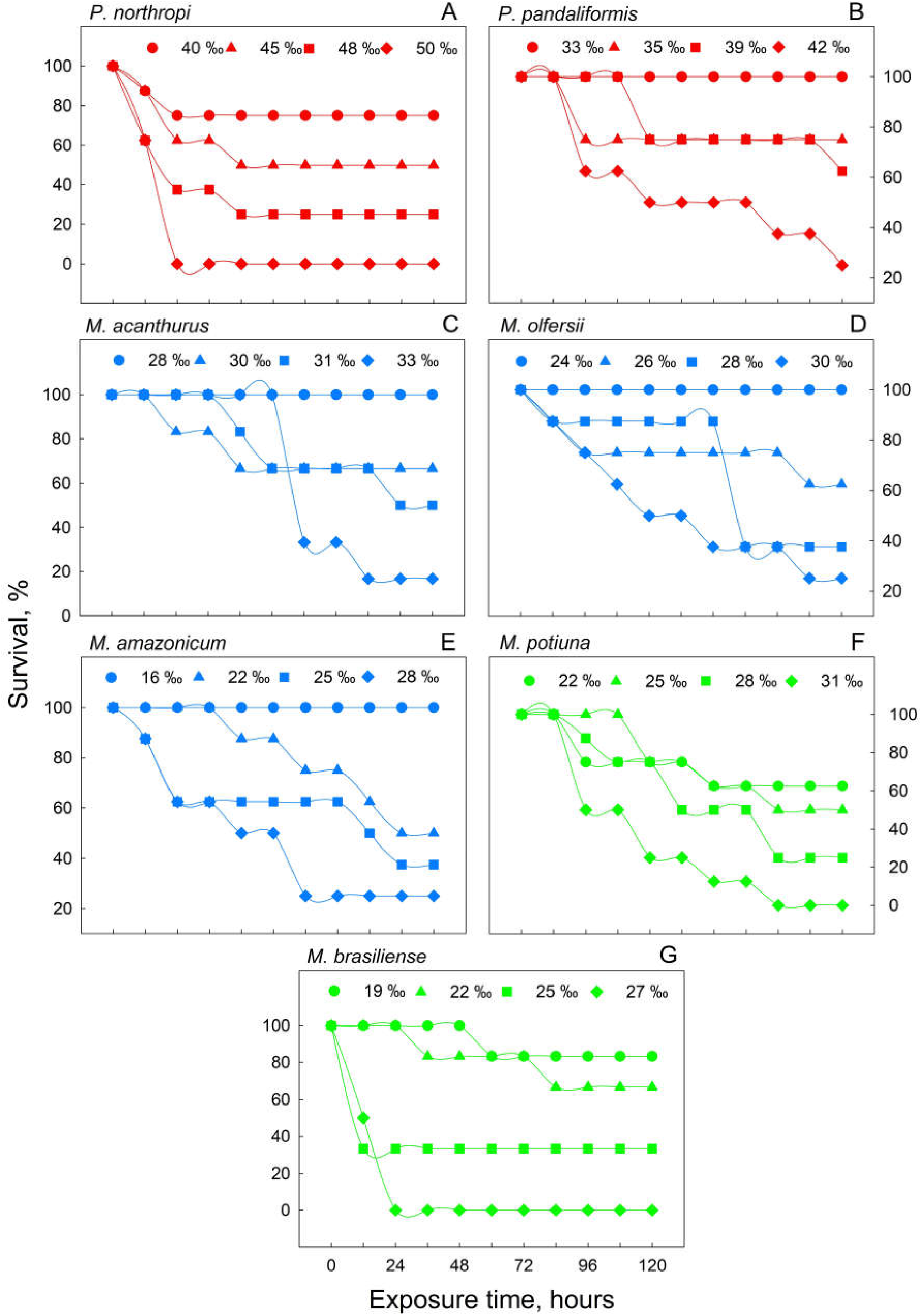
Survival curves for palaemonid shrimp species from different habitats exposed to a hyperosmotic salinity challenge for 5 days. A, *Palaemon northropi* a marine tide pool shrimp. B, *P. pandaliformis* an estuarine dweller. C, D and E, *Macrobrachium acanthurus, M. olfersii* and *M. amazonicum* (land-locked population) diadromous freshwater shrimps. F and G, *M. potiuna* and *M. brasiliense* hololimnetic freshwater shrimps. All species were subjected directly to four different salinities (‰S) chosen over different ranges to establish their upper lethal salinity limits (UL_50_) after 120 h exposure (N= 8). Exposure time = 0 h for *P. northropi* and *P. pandaliformis* corresponds to the laboratory acclimatization salinities of 18 and 17 ‰S, respectively; for the *Macrobrachium* species Exposure time = 0 h corresponds to fresh water (<0.5 ‰S).

### Hemolymph osmolality, chloride concentration and muscle water content

*Palaemon northropi* maintained its hemolymph hyperosmotic (≈620 mOsm/kg H_2_O) to the control acclimatization salinity (18 ‰S, 540 mOsm/kg H_2_O) while *P. pandaliformis* was roughly isosmotic (520 mOsm/kg H_2_O) (17 ‰S, 510 mOsm/kg H_2_O). On 24-h exposure to their 80%UL_50_ salinities (35.0 and 32.0 ‰S respectively), both species strongly hypo-regulated hemolymph osmolality (695 and 559 mOsm/kg H_2_O, Figure 2). Osmolality in *P. pandaliformis* increased transiently after 24 h exposure but declined to control values after 120 h. All the *Macrobrachium* species strongly hyper-regulated their hemolymph osmolality between 360 and 450 mOsm/kg H_2_O when in fresh water (Figure 2), osmolality increasing sharply, however, after 24 h exposure at their 80%UL_50_ salinities, and moderately after 120 h, most becoming isosmotic. *Macrobrachium acanthurus* alone remained hypo-osmotic (593 mOsm/kg H_2_O) after 120 h acclimation (25 ‰S, 750 mOsm/kg H_2_O).

**Figure 2.**
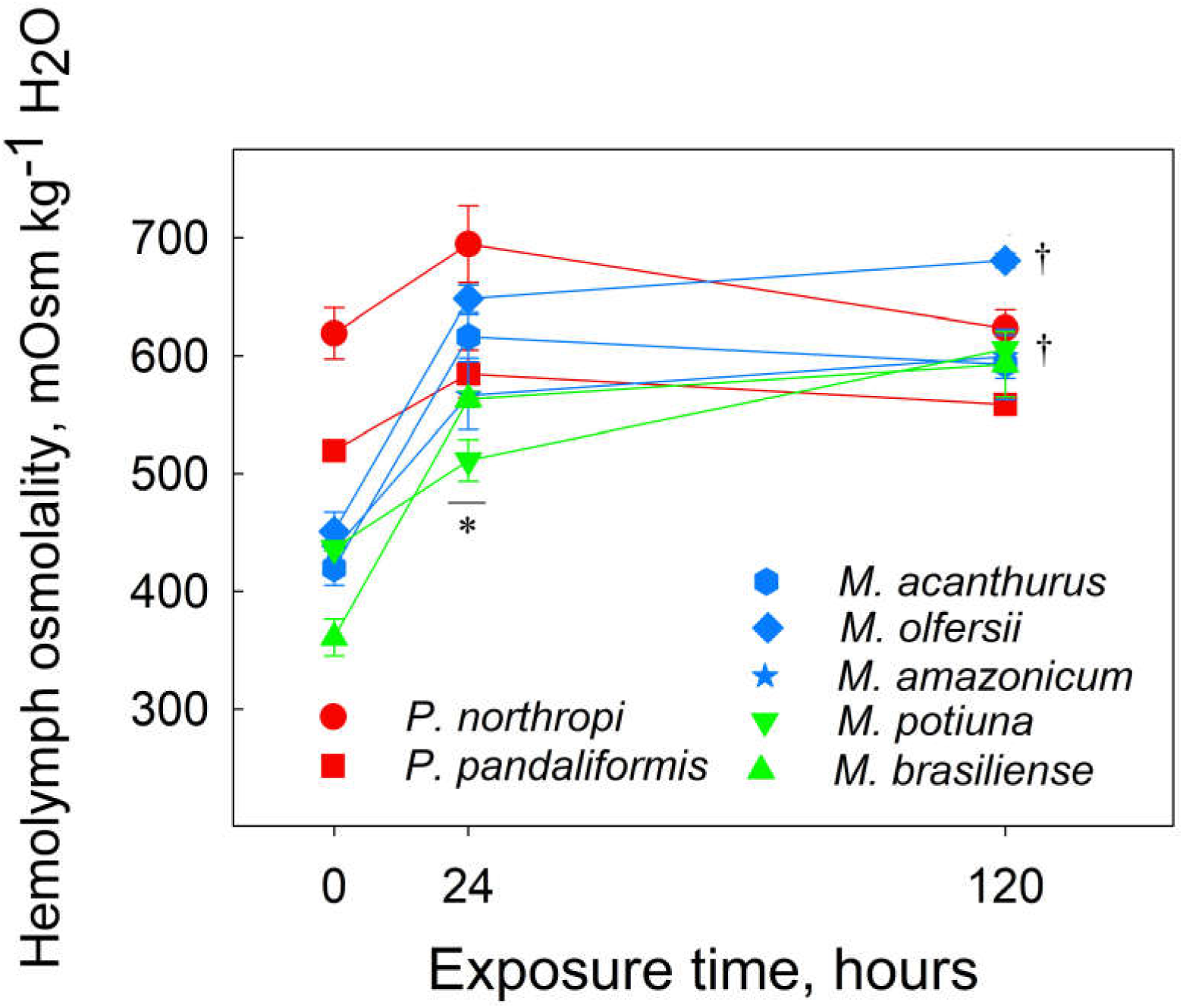
Hemolymph osmolalities of palaemonid shrimps from different habitats exposed to a hyperosmotic salinity challenge for 5 days. Shrimps were transferred directly from their control salinities (Time = 0 h) [18 ‰S, 540 mOsm/kg H_2_O for the tide pool shrimp, *Palaemon northropi*; 17 ‰S, 510 mOsm/kg H_2_O for the estuarine shrimp *P. pandaliformis*; and fresh water (<0.5 ‰S, <15 mOsm/kg H_2_O) for the diadromous and hololimnetic *Macrobrachium* species] to their respective 80%UL_50_ salinities (*P. northropi* 35 ‰S [1,050 mOsm kg^-1^ H_2_O], *P. pandaliformis* 32 [960 mOsm kg^-1^ H_2_O], *M. acanthurus* 25 [750 mOsm kg^-1^ H_2_O], *M. olfersii* 22 [660 mOsm kg^-1^ H_2_O], *M. amazonicum, M. potiuna* 19 [570 mOsm kg^-1^ H_2_O] and *M. brasiliense* 20 ‰S [600 mOsm kg^-1^ H_2_O]). Hemolymph from individual shrimps was sampled after 24 or 120 h. Data are the mean ± SEM (7≤N≤9). *P≤0.05 compared to control group (Time = 0 h) for all species except *P. northropi*. ^†^P=0.05 compared to 0 and 24 h for *M. olfersii* and *M. potiuna*.

Hemolymph Cl^-^ was unaltered by acclimation to the 80%UL_50_ salinities in the *Palaemon* species and was strongly hypo-regulated between 237 and 251 mmol Cl^-^ L^-1^ after 120 h (Figure 3). Despite increasing markedly by ≈1.6-fold to ≈250 mmol Cl^-^ L^-1^ after 24-h salinity exposure, hemolymph Cl^-^ in the *Macrobrachium* species, unlike osmolality, was strongly hypo-regulated during acclimation (≈240 mmol Cl^-^ L^-1^). There was little or no change after 120 h acclimation (Figure 3) except for *M. acanthurus* in which hemolymph Cl^-^ was notably hypo-regulated.

**Figure 3.**
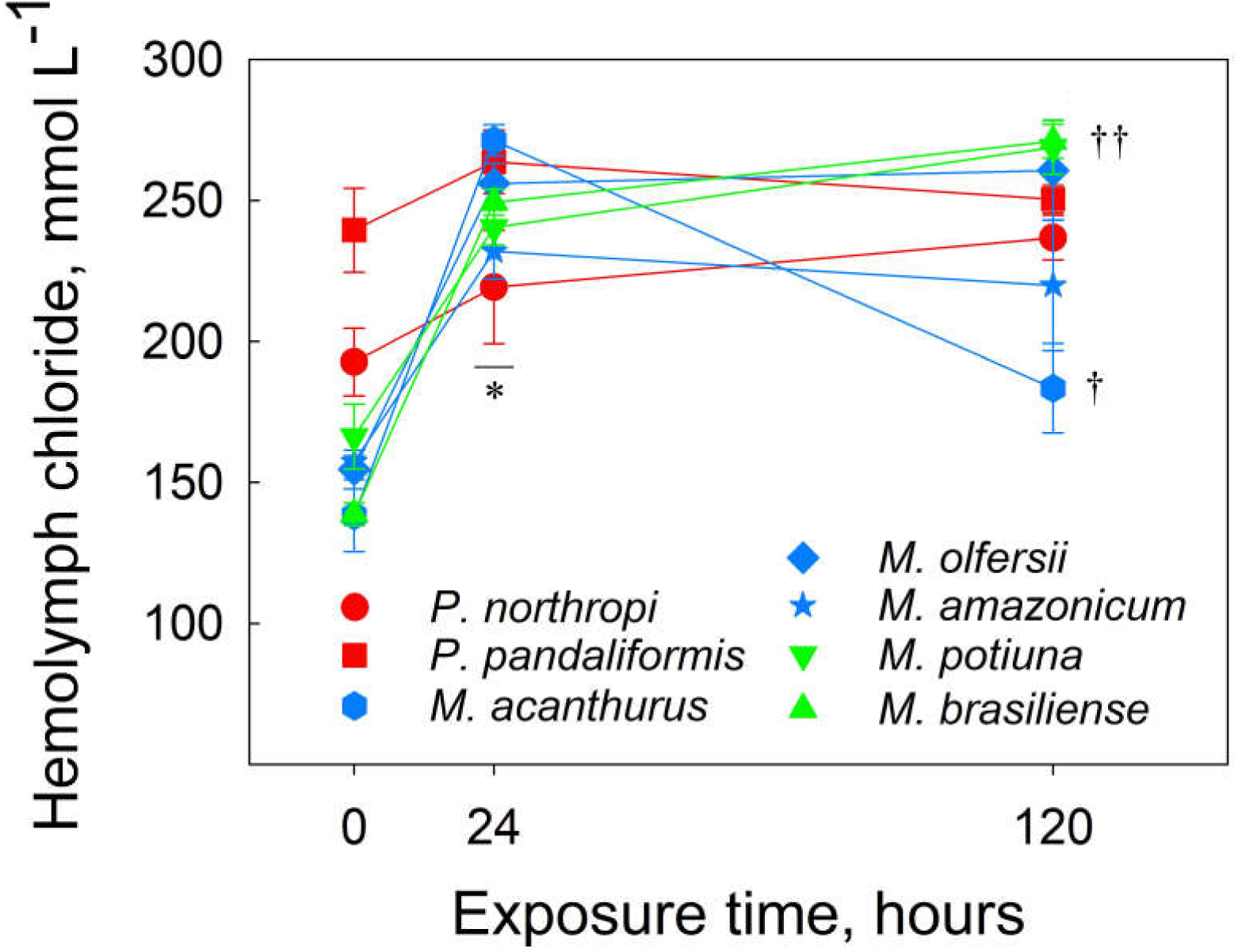
Hemolymph chloride concentrations in palaemonid shrimps from different habitats exposed to a hyperosmotic salinity challenge for 5 days. Shrimps were transferred directly from their control salinities (Time = 0 h) [18 ‰S, 288 mmol Cl^-^ L^-1^ for the tide pool shrimp, *Palaemon northropi*; 17 ‰S, 272 mmol Cl^-^ L^-1^ for the estuarine shrimp *P. pandaliformis*; and fresh water (<0.5 ‰S, <8 mmol Cl^-^ L^-1^) for the diadromous and hololimnetic *Macrobrachium* species] to their respective 80%UL_50_ salinities (*P. northropi* 35 ‰S [560 mmol Cl^-^ L^-1^], *P. pandaliformis* 32 [512 mmol Cl^-^ L^-1^], *M. acanthurus* 25 [400 mmol Cl^-^ L^-1^], *M. olfersii* 22 [352 mmol Cl^-^ L^-1^], *M. amazonicum* and *M. potiuna* 19 [304 mmol Cl^-^ L^-1^] and M. brasiliense 20 ‰S [320 mmol Cl^-^ L^-1^]). Hemolymph from individual shrimps was sampled after 24 or 120 h. Data are the mean ± SEM (7≤N≤9). *P≤0.05 compared to control group (Time = 0 h) for all species. ^†^P≤0.05 compared to 0 and 24 h for *M. acanthurus, M. potiuna* and *M. brasiliense*.

Muscle water content increased by ≈2% in *P. northropi* after 120-h exposure, and in *P. pandaliformis* diminished by ≈3% after 24 h but increased by ≈4.5% after 120 h (Figure 4). In *M. acanthurus, M. olfersii* and *M. amazonicum*, muscle water content decreased by ≈3% after 24 h, returning to control values after 120 h. Muscle water content was unchanged during acclimation in *M. potiuna* but decreased by ≈2% in *M. brasiliense*.

**Figure 4.**
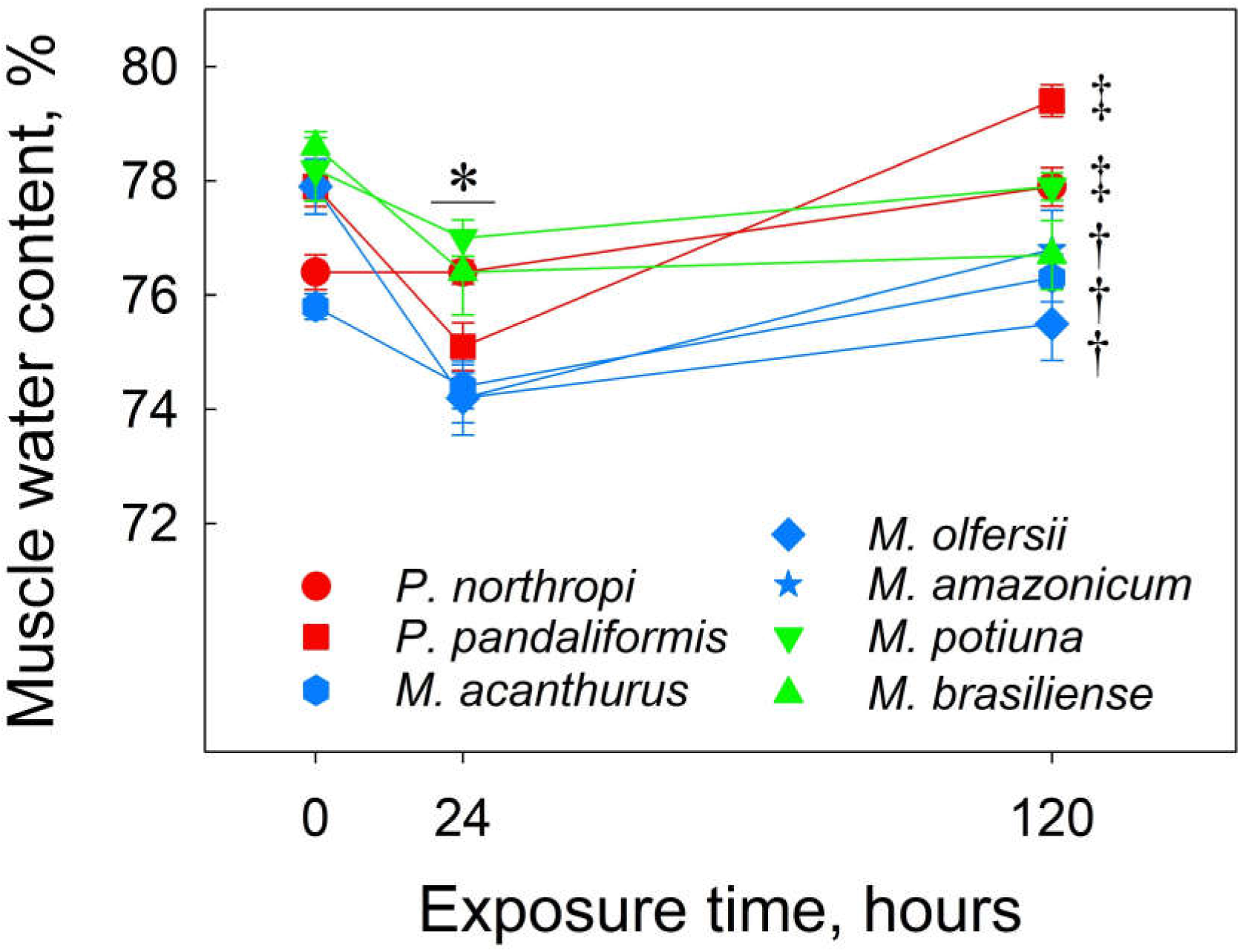
Muscle water content of palaemonid shrimps from different habitats exposed to a hyperosmotic salinity challenge for 5 days. Shrimps were transferred directly from their control salinities (Time = 0 h) [18 ‰S, 540 mOsm/kg H_2_O for the tide pool shrimp, *Palaemon northropi*; 17 ‰S, 510 mOsm/kg H_2_O for the estuarine shrimp *P. pandaliformis*; and fresh water (<0.5 ‰S, <15 mOsm/kg H_2_O) for the diadromous and hololimnetic *Macrobrachium* species] to their respective 80%UL_50_ salinities (*P. northropi* 35 ‰S, *P. pandaliformis* 32, *M. acanthurus* 25, *M. olfersii* 22, *M. amazonicum, M. potiuna* 19 and *M. brasiliense* 20 ‰S). Abdominal muscle tissue from individual shrimps was sampled after 24 or 120 h. Data are the mean ± SEM (7≤N≤9). *P≤0.05 compared to control group (Time = 0 h) for all species except *P. northropi* and *M. potiuna*. ^†^P≤0.05 compared to 24 h for *M. acanthurus, M. olfersii* and *M. amazonicum*. ^‡^P≤0.05 compared to 0 and 24 h for *P. northropi* and *P. pandaliformis*.

### Partial sequences of the gill sodium-potassium-two chloride symporter and ribosomal protein L10 genes

Partial sequences for the gill NKCC gene from the seven palaemonid shrimps were obtained after amplification and cloning, requiring two different primer pairs based on the complete NKCC sequence for *Macrobrachium koombooloomba*. Primer pair NKCC_Mk_F1/R1 produced a 415-bp product in *M. acanthurus, M. amazonicum, M. brasiliense* and *M. potiuna*, which were deposited with GenBank under accession numbers MG38514.0, MG38514.1, MG38514.2 and MG38514.3, respectively. For *P. northropi, P. pandaliformis* and *M. olfersii*, a 561-bp product was obtained using primer pair NKCC_Mk_F2/R2, deposited under Genbank accession numbers MG652471.1, MG652469.1 and MG652470.1, respectively.

Multiple alignments among the *Macrobrachium* nucleotide sequences (except *M. olfersii*) obtained using primer pair NKCC_Mk_F1/R1 revealed an elevated overall identity of ≈95%, ranging from 94.0% between *M. acanthurus* and *M. potiuna* to 96.9% between *M. acanthurus* and *M. amazonicum*. Alignment of the partial sequences for *P. northropi, P. pandaliformis* and *M. olfersii* obtained using primer pair NKCC_Mk_F2/R2 revealed much lower identities between the *Palaemon* species (46.6%), but reached 88.6% between *P. pandaliformis* and *M. olfersii*. Alignment of the deduced amino acid sequences among the *Macrobrachium* species (except *M. olfersii)* ranged from 94.6 to 96.7% identity (Figure 5), and from 92 to 92.9% among *P. northropi, P. pandaliformis* and *M. olfersii* (Figure 6). Amino acid identities of the *Palaemon* species and *M. olfersii* with the NKCC1 isoforms from the atyid shrimp *Halocaridina rubra* (AIM43576.1) were 82%, and 75% with the penaeid shrimp *Penaeus vannamei* (ROT83241.1). The amino acid sequences for *M. acanthurus, M. amazonicum, M. potiuna* and *M. brasiliense* showed 70 to 75% identity with the *H. rubra* isoform.

**Figure 5.**
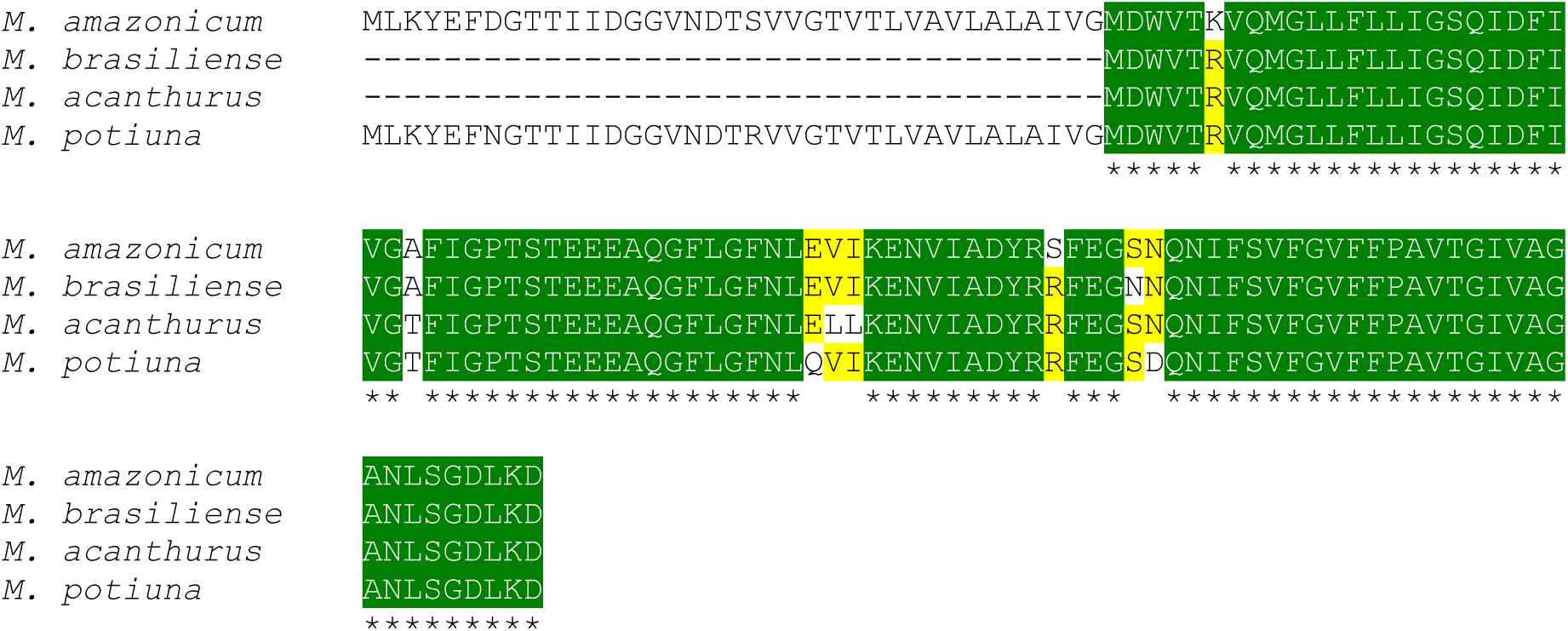
Multiple alignments of deduced amino acid sequences for the gill sodium-potassium-two chloride symporter in several species of freshwater shrimps (*Macrobrachium*). Alignment was performed using the Clustal Omega software package (http://www.ebi.ac.uk/Tools/msa/clustalo). Identical amino acids among all four species are shown in green (*) (100% identity) with three identical residues in yellow (75% identity). A alanine, C cysteine, D aspartate, E glutamate, F phenylalanine, G glycine, H histidine, I isoleucine, K lysine, L leucine, M methionine, N asparagine, P proline, Q glutamine, R arginine, S serine, T threonine, V valine, W tryptophan, Y tyrosine.

**Figure 6.**
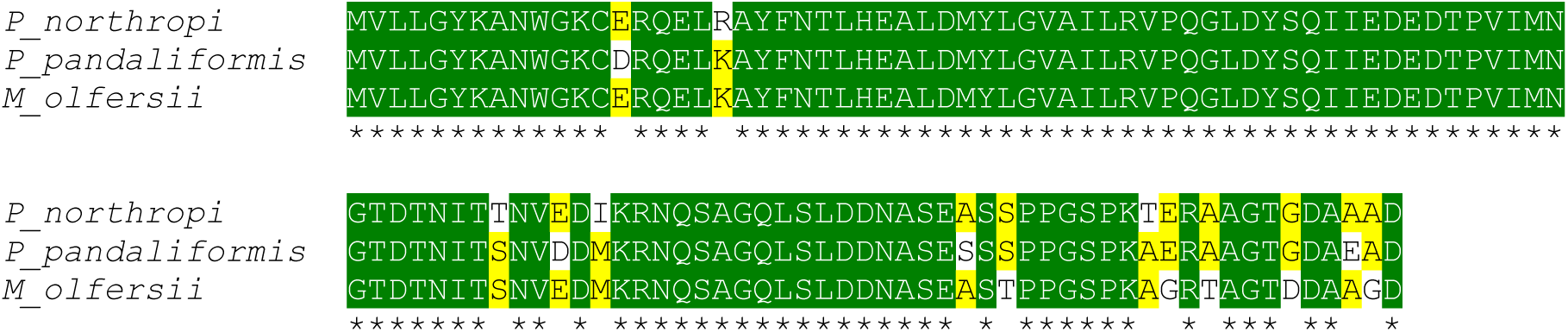
Multiple alignments of deduced amino acid sequences for the gill sodium-potassium-two chloride symporter in three species of palaemonid shrimps (*Palaemon* and *Macrobrachium*). Alignment was performed using the Clustal Omega software package (http://www.ebi.ac.uk/Tools/msa/clustalo). Identical amino acids among all three species are shown in green (*) (100% identity) with two identical residues in yellow (66.7% identity). A alanine, C cysteine, D aspartate, E glutamate, F phenylalanine, G glycine, H histidine, I isoleucine, K lysine, L leucine, M methionine, N asparagine, P proline, Q glutamine, R arginine, S serine, T threonine, V valine, W tryptophan, Y tyrosine.

Partial nucleotide sequences for the gill RPL10 gene from the seven palaemonid shrimps were obtained after amplification and cloning. The RPL10_Cs_F/R primer pair produced a 251-bp fragment for the gill RPL10 gene that was sequenced in *P. pandaliformis* and *M. potiuna* and deposited with GenBank under accession numbers KP890671.1 and KU726244.1, respectively. The partial sequences for the other species were obtained previously by us using the same primer pair (RPL10_Cs_F/R), and available from Genbank (*P. northropi* JN251135, *M. acanthurus* JN251134, *M. amazonicum* GU366065 and *M. brasiliense* JN251133).

Multiple alignment of these RPL10 nucleotide sequences showed the highest identity between *M. acanthurus* and *M. amazonicum* (99.6%). Other comparisons among *Macrobrachium* species ranged from 96 to 97.2%. The lowest sequence identities were found between *P. pandaliformis* and the other species (88.8 to 90.8%). The respective, predicted, partial amino acid sequences for the gill RPL10 are given in Figure 7. Amino acid identities among all species were 100%, except for *M. brasiliense* that bears a single different amino acid (87.5%).

**Figure 7.**
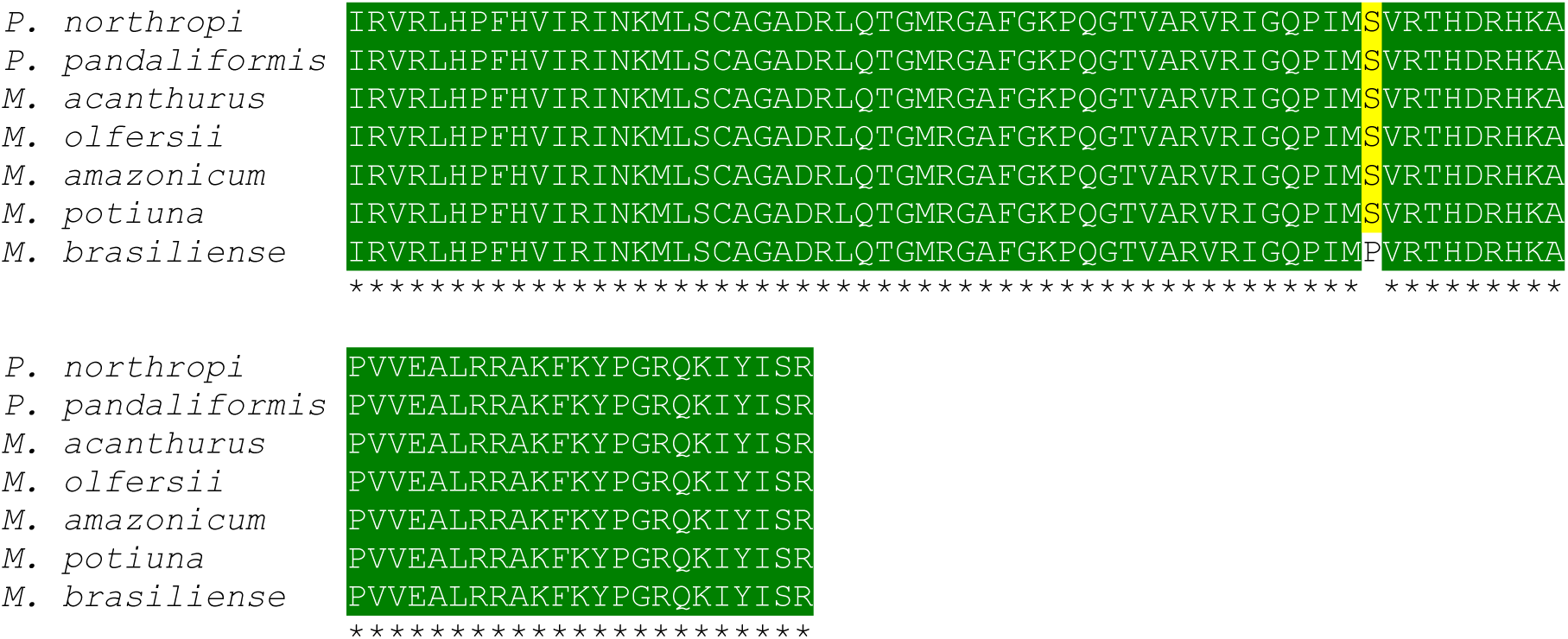
Multiple alignments of deduced amino acid sequences for the gill ribosomal protein L10 in seven species of palaemonid shrimp (Macrobrachium and Palaemon). Alignment was performed using the Clustal Omega software package (http://www.ebi.ac.uk/Tools/msa/clustalo). Identical amino acids among all seven species are shown in green (*) (100% identity) with six identical residues in yellow (85.7% identity). A alanine, C cysteine, D aspartate, E glutamate, F phenylalanine, G glycine, H histidine, I isoleucine, K lysine, L leucine, M methionine, N asparagine, P proline, Q glutamine, R arginine, S serine, T threonine, V valine, W tryptophan, Y tyrosine.

### mRNA expression and protein abundance of the gill sodium-potassium-two chloride symporter

The relative mRNA expression of the gill sodium-potassium-two chloride symporter was unaffected by 80%UL_50_ salinity challenge (0.198<F<3.09, 0.083<P<0.823) (Figure 8), although expression in *M. amazonicum* and *M. potiuna* tended to increase by ≈1.4-fold after 120 h. Basal expression was greatest in *P. pandaliformis*, reaching 240-fold that for *M. olfersii* and 660-fold that of *M. amazonicum*, the lowest (both P<0.001) (Figure 8). Even after 120 h hyper-osmotic challenge, expression in *P. pandaliformis* remained elevated, ≈450-fold that of *M. brasiliense*.

**Figure 8.**
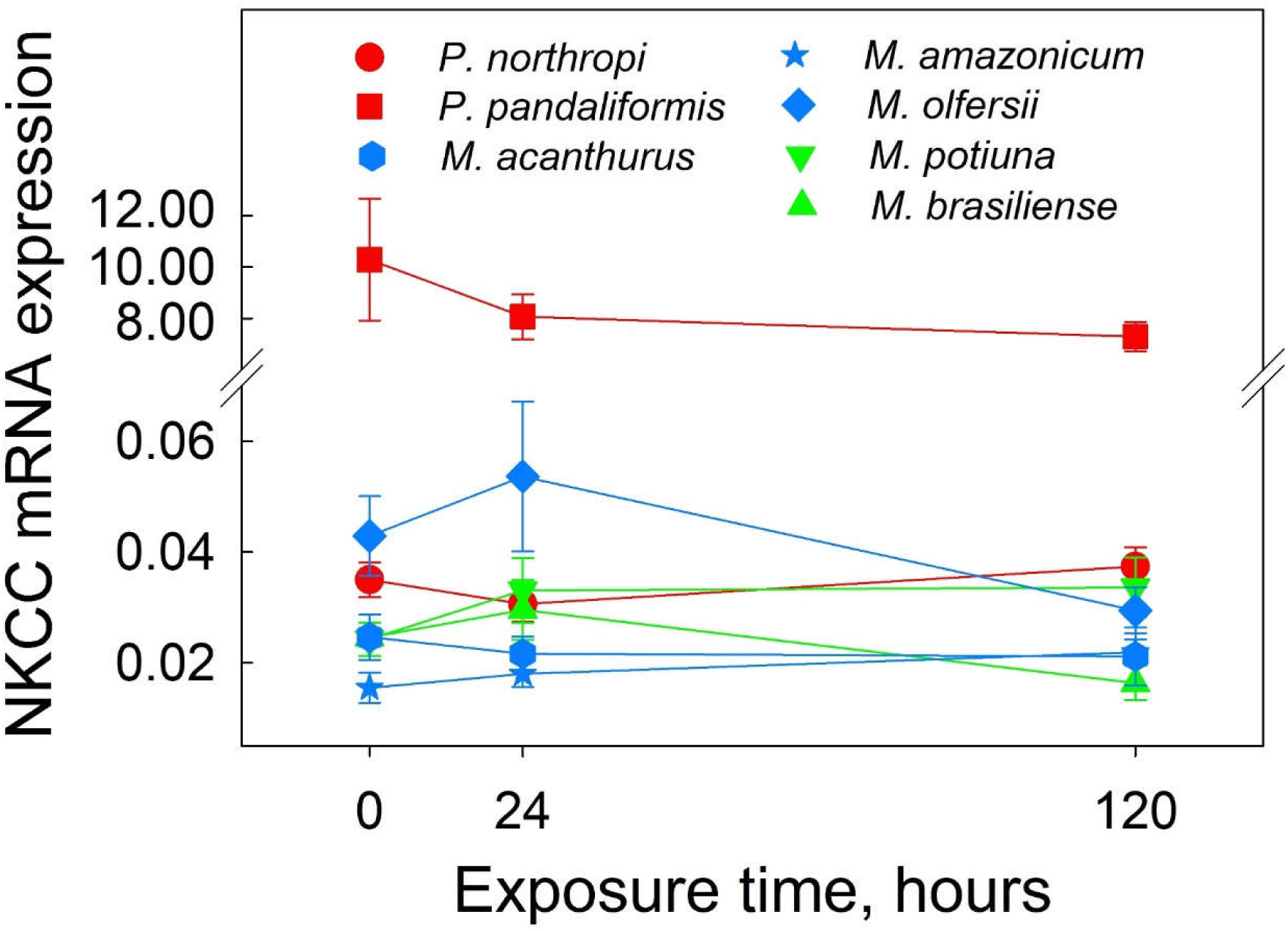
Relative gene expression of the gill sodium-potassium-two chloride symporter (NKCC) in several species of palaemonid shrimps (*Palaemon* and *Macrobrachium*) from different salinity habitats exposed to a hyperosmotic salinity challenge (80%UL_50_) for 5 days. Exposure time to the different salinities had no effect on NKCC gene expression. Shrimps were transferred directly from their control acclimatization salinities (Time = 0 h) [18 ‰S, 540 mOsm/kg H_2_O for the tide pool shrimp, *Palaemon northropi*; 17 ‰S, 510 mOsm/kg H_2_O for the estuarine shrimp *P. pandaliformis*; and fresh water (<0.5 ‰S, <15 mOsm/kg H_2_O) for the diadromous and hololimnetic *Macrobrachium* species] to their respective 80%UL_50_ salinities (*P. northropi* 35 ‰S, *P. pandaliformis* 32, *M. acanthurus* 25, *M. olfersii* 22, *M. amazonicum, M. potiuna* 19 and *M. brasiliense* 20 ‰S). Gills from individual shrimps was sampled after 24 or 120 h. NKCC expressions (mean ± SEM, N=8) were calculated using the comparative CT method (2^-ΔΔCT^) and have been normalized by expression of the constitutive gene RPL10 in the same sample. No significant differences were detected (P>0.05).

In contrast, protein expression of the gill sodium-potassium-two chloride symporter showed clear genus- and species-specific responses to 80%UL_50_ salinity challenge (Figure 9). Expression was unchanged in the two marine-brackish water *Palaemon* species and in the coastal hololimnetic freshwater species, *M. potiuna* (0.316<F<1.57, 0.236<P<0.734). In the diadromous species, expression increased 2.2-(F=4.12, P=0.044), 2.1-(F=3.91, P=0.043) and 1.7-fold (F=5.67, P=0.017) respectively in *M. acanthurus, M. amazonicum* and *M. olfersii* after 120 h acclimation. In the continental hololimnetic species, *M. brasiliense*, expression increased 1.6-fold after 24 h and 2.2-fold after 120 h exposure (F =7.03, P=0.007) (Figure 9).

**Figure 9.**
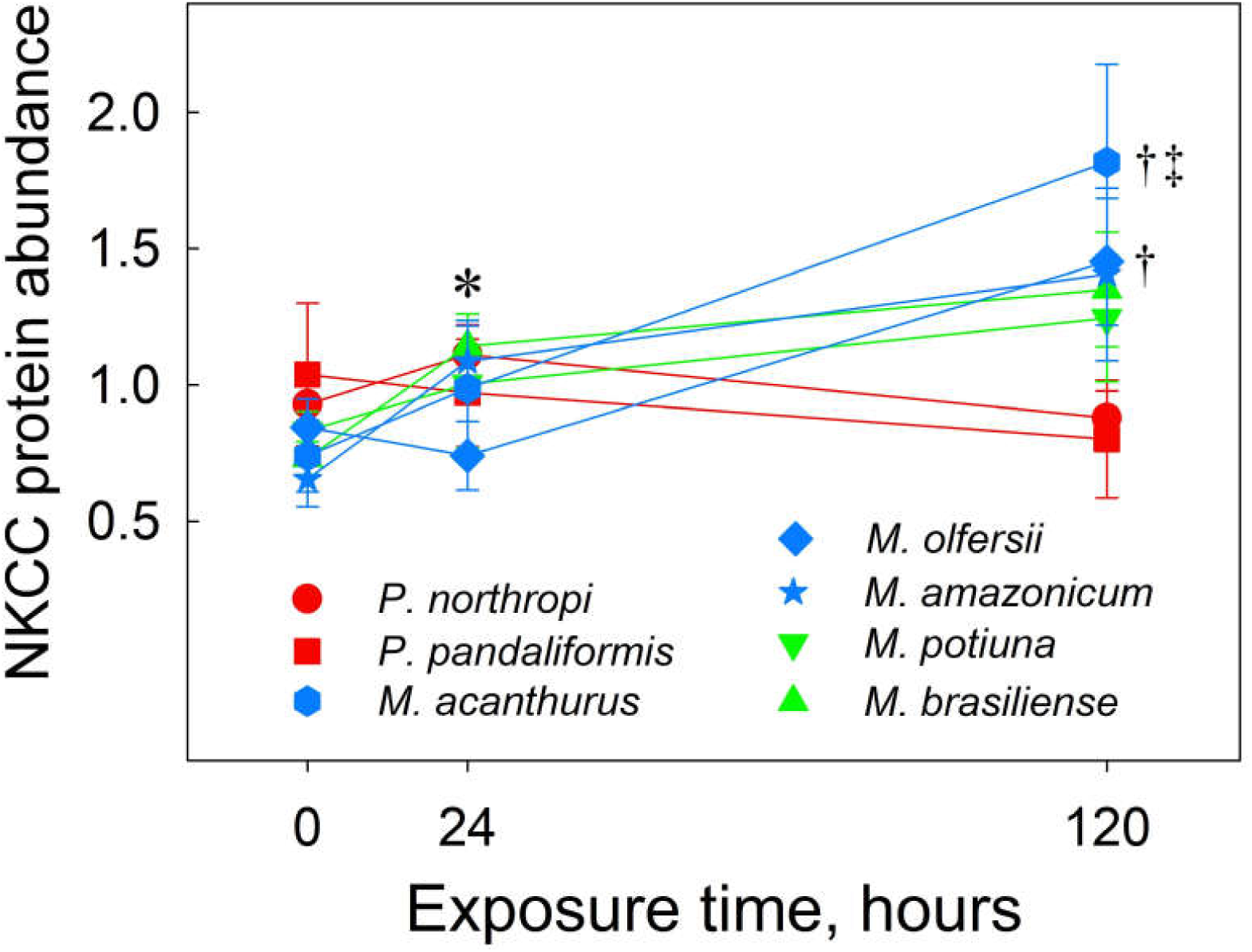
Protein expression of the gill sodium-potassium-two chloride symporter (NKCC) in several species of palaemonid shrimps (*Macrobrachium* and *Palaemon*) from different salinity habitats exposed to hyperosmotic salinity challenge (80%UL_50_) for 5 days. Abundance is unchanged in the *Palaemon* species but increases with exposure time in the diadromous and hololimnetic *Macrobrachium* species. Shrimps were transferred directly from their control acclimatization salinities (Time = 0 h) [18 ‰S, 540 mOsm/kg H_2_O for the tide pool shrimp, *Palaemon northropi*; 17 ‰S, 510 mOsm/kg H_2_O for the estuarine shrimp *P. pandaliformis*; and fresh water (<0.5 ‰S, <15 mOsm/kg H_2_O) for the diadromous and hololimnetic *Macrobrachium* species] to their respective 80%UL_50_ salinities (*P. northropi* 35 ‰S, *P. pandaliformis* 32, *M. acanthurus* 25, *M. olfersii* 22, *M. amazonicum, M. potiuna* 19 and *M. brasiliense* 20 ‰S). Gills from individual shrimps were sampled after 24 or 120 h. NKCC protein expressions (mean ± SEM, N=8) are given as the ratio of NKCC expression to that of the reference protein, β-actin, in the same sample. *P<0.05 compared to control group (Time = 0 h) for *M. brasiliense*. ^†^P≤0.05 compared to 0 and 24 h for *M. acanthurus* and *M. olfersii*. ^‡^P≤0.05 compared to control group (Time = 0 h) for *M. amazonicum*.

### Phylogenetic signal, correlations and comparisons

UL_50_, hemolymph osmolality and Cl^-^ concentration in the control acclimatization salinities, and Cl^-^ hypo-regulation and protein expression of the gill sodium-potassium two chloride symporter after 120 h 80%UL_50_ salinity challenge showed positive autocorrelations (Moran’s *I* >0.71, P<0.05), demonstrating strong phylogenetic structuring. Gene expression of the sodium-potassium-two chloride symporter under all conditions, protein expression in the control acclimatization salinities and at the 80%UL_50_ after 24 h, and muscle water content and hemolymph osmolality at the 80%UL_50_ salinity were not phylogenetically structured, which reveals that interspecific variability in this case may have been driven by abiotic factors like salinity.

Overall, there was no correlation between the sodium-potassium-two chloride symporter gene or protein expression and the osmoregulatory parameters (i. e., hemolymph osmolality and Cl^-^ concentration) for any species under any experimental condition (0.07<F< 2.88, 0.15<P< 0.81). However, gene expression and hemolymph [Cl^-^] were correlated in the control acclimatization salinities (F=13.9, P=0.014). Phylogenetic paired t-testing revealed no difference between the relative mRNA expression at the 80%UL_50_ after 24 h (T = −0.525, P = 0.628) and 120 h (T = 1.296, P = 0.265) compared to the control acclimatization group (Time = 0 h) for any species.

However, protein expression of the sodium-potassium-two chloride symporter tended to correlate with hemolymph osmolality in the control acclimatization salinity (F=5.249; P=0.071), and there was an increase in expression after 24 h compared to Time = 0 h (T = −3.382, P = 0.028).

## DISCUSSION

We have characterized the survival and osmoregulatory abilities of several palaemonid shrimps from different osmotic niches when challenged by elevated salinity for 5 days. The *Palaemon* species were more tolerant and were excellent hypo-osmotic and -chloride regulators, followed by the diadromous *Macrobrachium* species. Gill NKCC mRNA transcription was unaltered on salinity challenge, despite the increase in gill NKCC protein synthesis, which was exclusive to the *Macrobrachium* species, except for the *M. potiuna*. The capability to *hyper*-regulate hemolymph [Cl^-^] correlated with gill NKCC symporter gene expression, while NKCC protein synthesis seems to be associated with *hyper*-osmoregulatory ability.

### Hyperosmotic challenge and survival

Upper salinity tolerances (UL_50_) showed phylogenetic signal and were highest in the *Palaemon* species, *i. e*., *P. northropi*, a marine tide pool shrimp, and *P. pandaliformis*, an estuarine dweller (43.1 and 39.7 ‰S, respectively). Other *Palaemon* species like *P. ritteri* (UL_50_ = 47.5 ‰S, Reynolds 1975), and *P. affinis* (>75% survival in 43 ‰S for 5 days, Kirkpatrick and Jones 1985), also from tide pools, or estuaries and mangroves, exhibit notable salinity tolerances. In contrast, the diadromous, freshwater *Macrobrachium* species, *M. acanthurus* and *M. olfersii*, collected from coastal rivers, exhibited intermediate UL_50_’s of 31.4 and 28.0 ‰S, respectively. The land-locked *M. amazonicum* population (24.5 ‰S), uninfluenced by brackish water, was similar to the hololimnetic species *M. potiuna* (24 ‰S) and *M. brasiliense* (24.7 ‰S).

*Macrobrachium* is a genus of mostly freshwater species. Of these, however, some like the Indo-Pacific *M. equidens*, inhabit brackish water environments, showing an upper salinity tolerance of ≈40 ‰S (Denne 1968), much higher than the hololimnetic *M. australiense* (UL_50_ of 25 ‰S) (Denne 1968) and the similar Brazilian species. Such high salinity tolerances (>25 ‰S), although variable within the genus, have been retained in species that do not encounter brackish water during their life cycle, which, corroborated by the significant phylogenetic signal for UL_50_, suggests that during the adaptive radiation of the group into fresh water, tolerance to high salinities has been conserved.

*Palaemon* species, even more tolerant of higher salinities, are also strong hyper/hypo-osmoregulators (Freire et al. 2003, Augusto et al. 2009, Gonzalez-Ortegón et al. 2015, Faleiros et al. 2017). *Palaemon northropi* and *P. pandaliformis* showed greater regulatory capabilities compared to *Macrobrachium* species like *M. olfersii, M. potiuna* and *M. brasiliense* (Freire et al. 2003, Freire et al. 2008b). Although *Palaemon* species can hyper-regulate hemolymph osmolality in dilute media (>2 ‰S, Kirkpatrick and Jones 1985, Freire et al. 2003, Foster et al. 2010, Faleiros et al. 2017), they show poor survival. *Palaemon ritteri* has a lower lethal salinity limit (LL_50_) of 10 ‰S (Reynolds 1975) while *P. pandaliformis* and *P. northropi* survive less than 3 h in fresh water (<0.5 ‰S) (Freire et al. 2003), the latter showing less than 45% survival in 1 ‰S (Augusto et al. 2009). *Palaemon affinis* exhibits less than 35% survival in 0.5 ‰S (Kirkpatrick and Jones 1985).

The survival of these palaemonid shrimps in saline media is phylogenetically structured, suggesting that osmotic tolerance has followed the cladogenetic evolution of the group during the course of occupation of distinct osmotic niches.

### Osmotic and ionic regulatory capability and tissue water content

The ability to hypo-regulate hemolymph osmolality showed phylogenetic signal, with closely related species exhibiting similar responses. The strong hypo-regulatory capability of *P. northropi* is sustained even at salinities well above those encountered in its natural tide pool habitat (3-35 ‰S, 90-1,050 mOsm/kg H_2_O). When acclimated to salinities up to 50 ‰S (1,500 mOsm/kg H_2_O) for 10 days, *P. northropi* holds its hemolymph osmolality at ≈1,150 mOsm/kg H_2_O, considerably below the external medium (Δ = −350 mOsm/kg H_2_O) (Faleiros et al. 2017). In seawater at 35 ‰S (1,050 mOsm/kg H_2_O, 80%UL_50_), *P. northropi* maintains its hemolymph osmolality at 624 mOsm/kg H_2_O after 5 days (Δ = −426 mOsm/kg H_2_O). *Palaemon pandaliformis* held at 32 ‰S (960 mOsm/kg H_2_O, 80%UL_50_) for 5 days also strongly hypo-regulates its hemolymph osmolality at 559 mOsm/kg H_2_O (Δ = −401 mOsm/kg H_2_O), and at ≈618 mOsm/kg H_2_O when in 35 ‰S for 17 h (Foster et al. 2010).

In contrast, *Macrobrachium* species exhibit strong hyper-osmoregulatory capabilities in fresh water, and show variable hypo-regulatory capacities at higher salinities (Moreira et al. 1983, Freire et al. 2003, Foster et al. 2010, Maraschi et al. 2015). To illustrate, *M. acanthurus* exhibited good hypo-regulatory ability (Δ= −157 mOsm/kg H_2_O) while *M. olfersii, M. amazonicum, M. brasiliense* and *M. potiuna* showed little hypo-osmoregulatory capacity after acclimation at their respective 80%UL_50_’s (Δ ≅0 mOsm/kg H_2_O). Curiously, the loss of hemolymph hypo-osmoregulatory ability was not accompanied by a loss in hemolymph Cl^-^ hypo-regulatory capacity.

Like hemolymph osmolality, the ability to hypo-regulate hemolymph Cl^-^ also showed phylogenetic signal consequent to the phylogenetic proximity between species. On acclimation, *Palaemon pandaliformis* maintained a smaller Cl^-^ gradient (Δ= −261) than did *P. northropi* (Δ= −323) at their respective 80%UL_50_’s of 32 and 35 ‰S. This reduction in gradient is even more evident in the *Macrobrachium* species. *Macrobrachium acanthurus* maintained a Cl^-^ gradient of −217 mM Cl^-^, whereas *M. olfersii* and *M. amazonicum* maintained gradients of Δ= −91 and Δ= −100 mM Cl^-^, respectively, and *M. brasiliense* and *M. potiuna* of just Δ= −49 and Δ= −35 mM Cl^-^, respectively, after 5 days exposure at their 80%UL_50_’s. Nevertheless, some ability to hypo-regulate hemolymph Cl^-^ at high salinity is preserved in all the palaemonids examined here. The phylogenetic signal noted for hemolymph osmolality and [Cl^-^] reveals that the phylogenetic component dominates inter-specific variability in these traits, which likely have been inherited from a weakly hyper-osmoregulating, brackish water (17 ‰S) palaemonid ancestor (McNamara and Faria 2012).

Most species, except *M. potiuna*, showed muscle water loss and variable recovery in response to acclimation at their respective 80%UL_50_’s. Increased or decreased osmolality and ion concentrations in the hemolymph result in the passive influx or efflux of cell water and ions and, thus, reduced tissue hydration is usually associated with an increase in the concentration of the internal *milieu* (Freire et al. 2008b, Freire et al. 2013, Maraschi et al. 2015, Freire et al. 2018). Compensatory mechanisms of isosmotic intracellular regulation (IIR) ensure cellular viability and are prerequisites for the invasion of osmotically distinct environments (Florkin and Schoffeniels 1969, Freire et al. 2008b). Palaemonid shrimps that have invaded the strictly freshwater environment as hololimnetic species have maintained a high capability for IIR (Freire et al. 2008b). Muscle tissue hydration in *M. potiuna* was unaffected by salinity challenge, despite the increases in hemolymph osmolality and [Cl^-^], which corroborates the species’ excellent IIR capability.

### Expression of the sodium-potassium-two chloride symporter

The capability for osmotic and ionic hypo-regulation in the Palaemonidae, even in strictly freshwater species, involves the regulation of genes responsible for downstream transepithelial ion transport (Towle and Weihrauch 2001, Faleiros et al. 2010, Havird et al. 2014, 2016, Rahi et al. 2017, 2019). Alterations in water permeability and ion fluxes across the gill epithelia, the rapid modulation of membrane ion transporting ATPases and intracellular enzyme activities, together with up-regulated gene transcription and protein synthesis, constitute some of the osmotically sensitive response mechanisms to salinity challenge (Roy and Bhoite 2015). Among the possible mechanisms underlying ion secretion, transport mediated by the electron neutral sodium-potassium-two chloride symporter plays a significant role in Cl^-^ secretion by the fish gill (Evans 2010).

The NKCC gene is likely involved in osmoregulation by palaemonid shrimps such as *Macrobrachium rosenbergii* (Barman et al. 2012), *M. australiense* (Moshtaghi et al. 2016, 2018) and *M. koombooloomba* (Rahi et al. 2017). In the freshwater species *M. australiense*, gill NKCC mRNA expression increases 1.7-fold and 2-fold after 24 h exposure to 5 or 10 ‰S, respectively, (Moshtaghi et al. 2018). Transcriptional regulation of the gill NKCC gene has been explored in crabs like *Neohelice granulata*, (Luquet et al. 2005), *Carcinus maenas* (Towle et al. 2011), *Callinectes sapidus*, (Havird et al. 2016) and *Portunus trituberculatus* (Lv et al. 2016), and in the atyid shrimp *Halocaridina rubra* (Havird et al. 2014). Expression increases 60-fold in *Neohelice granulata* (Luquet et al. 2005) and 9.6-fold in *P. trituberculatus* (Lv et al. 2016) on exposure to 45 ‰S. Similarly, gill NKCC1 mRNA expression in the European yellow eel (Cutler and Cramb 2002), sea bass (Lorin-Nebel et al. 2006) and Atlantic salmon (Mackie et al. 2007) are higher when in seawater compared to fish acclimated to fresh water.

The evolutionary history of Cl^-^ regulation has not been explored in crustaceans. Gill NKCC gene expression and hemolymph Cl^-^ *hyper*-regulation were significantly correlated (P=0.014) in the *Palaemon* and *Macrobrachium* species examined here. However, there was no correlation between acclimation at the 80%UL_50_’s and gill NKCC gene expression, and no difference in NKCC expression after 80%UL_50_ challenge for 24 and 120 h compared to acclimatized controls. Thus, contrary to our hypothesis, the ability to secrete Cl^-^ is independent of augmented NKCC gene expression.

Gene expression is highly plastic and can be regulated by external factors (Wray 2013). The lack of response of the gill NKCC gene expression to salinity acclimation suggests that expression already may be maintained at a high level, as seen in *P. pandaliformis*, and even in *Macrobrachium* species in fresh water, or near isochloremicity in *Palaemon*, since the symporter also plays a role in salt uptake. Gill NKCC gene expression is up-regulated in the euryhaline shrimp *Halocaridina rubra* at 2 ‰S (Havird et al. 2014) and in the mud crab *Scylla paramamosain* in the post-molt, ion-uptake period (Xu et al. 2017).

Protein synthesis of the gill NKCC symporter increased up to 2-fold on salinity acclimation in the *Macrobrachium* species, particularly in the diadromous freshwater species *M. acanthurus, M. olfersii* and *M. amazonicum* that share dependence on brackish water for complete larval development. In the hololimnetic *M. brasiliense*, NKCC protein expression increased 1.5-fold. Phylogenetic t-testing revealed a transient increase in gill NKCC protein expression after 24 h at the 80%UL_50_ compared to acclimatized controls, declining after 120 h, which suggests that up-regulated NKCC protein expression results from post-translational events since gene expression is unaltered. A similar response occurs in Atlantic salmon after transfer to seawater (Pelis et al. 2001, Tipsmark et al. 2002). Inhibition of the NKCC symporter by furosemide in several decapod species reveals its participation in the regulation of intracellular volume and tissue hydration, in addition to a putative role in cellular Cl^-^ uptake and efflux (Freire et al. 2013).

Increased NKCC symporter gene translation in response to salinity challenge appears to have evolved as a cladogenetic event within the Palaemonidae and has been preserved among some extant *Macrobrachium* species (Barman et al. 2012, Moshtaghi et al. 2016). However, the 2-fold increase over ambient salinity to which *P. northropi* and *P. pandaliformis* were acclimated may have been insufficient to elicit a significant alteration in NKCC protein synthesis. These two palaemonids confront a mean daily salinity range of between 3 and ≈35 ‰S as a consequence of rainfall or evaporation. Many factors influence the rate of Cl^-^ transport mediated by the NKCC symporter, *e. g*., cell volume reduction, chemical messengers, reduced oxygen tension, increased intracellular [Mg^2+^], and reduced [Cl^-^] (reviewed in Flatman 2002), mediated via phosphorylation of protein domains, protein-protein interactions and the direct effect of cellular [Cl^-^] (Evans et al. 2010, Yuan et al. 2017). Rapid regulatory mechanisms, acting at levels other than transcription and translation, may provide an energy-efficient alternative for species confronting natural variations in salinity, such as seen in *Palaemon* species.

The hypo-osmotic and ionic regulatory mechanisms of palaemonid shrimps are thought to be similar to those of other hypo-osmoregulating crustaceans and marine vertebrates in general (McNamara and Faria 2012, Larsen et al. 2014). However, electrophysiological data show that the transepithelial voltage of crustaceans is externally positive, opposing that expected from a mechanism similar to that of vertebrates (reviewed in Larsen et al. 2014). Clearly, electrophysiological analyses employing the appropriate channel inhibitors, including measurements of real-time Cl^-^ fluxes and a search for active chloride-dependent ATPases are necessary to better elucidate this issue.

This novel characterization of the sodium-potassium-two chloride transporter at the transcriptional and translational levels in palaemonid shrimps from various osmotic niches, together with survival ability and measurements of hemolymph osmolality and Cl^-^ concentration, partially elucidates the as yet obscure mechanism of Cl^-^ hypo-regulation in palaemonids. The inclusion of phylogenetic relationships among the species evaluated here provides an adaptive interpretation of the role of the NKCC symporter in ion absorption and secretion, in addition to revealing a role for shared ancestry in Cl^-^ regulation in the Palaemonidae.

## ACKNOWLEDGMENTS

Shrimp collections were authorized under SISBIO permit #29594-12 to JCM issued by the Brazilian Ministério do Meio Ambiente, Instituto Chico Mendes de Conservação da Biodiversidade. We are grateful to Drs Rogério Faleiros and Mariana Capparelli for assistance with fieldwork. We thank Prof. Carolina A. Freire and Dr Viviane Prodocimo (Departamento de Fisiologia, UFPA) for laboratory support and Susie Teixeira Keiko for technical assistance. We are indebted to Dr Ademilson Panunto Castelo for access to the CFX96 Real-Time PCR Detection System. This investigation is part of a Ph D thesis submitted by ACM to the Graduate Program in Comparative Biology, Departamento de Biologia, FFCLRP/USP.

## Ethics approval

All institutional and federal guidelines followed, Environmental permit permit #29594-12 to JCM

## Consent to participate

Not applicable.

## Consent for publication

All authors read the manuscript and approve submission.

## Availability of data and material

Gene sequences deposited with NCBI GenBank.

## Code availability

Not applicable.

## Supplementary material

**Supplementary Figure 1.**
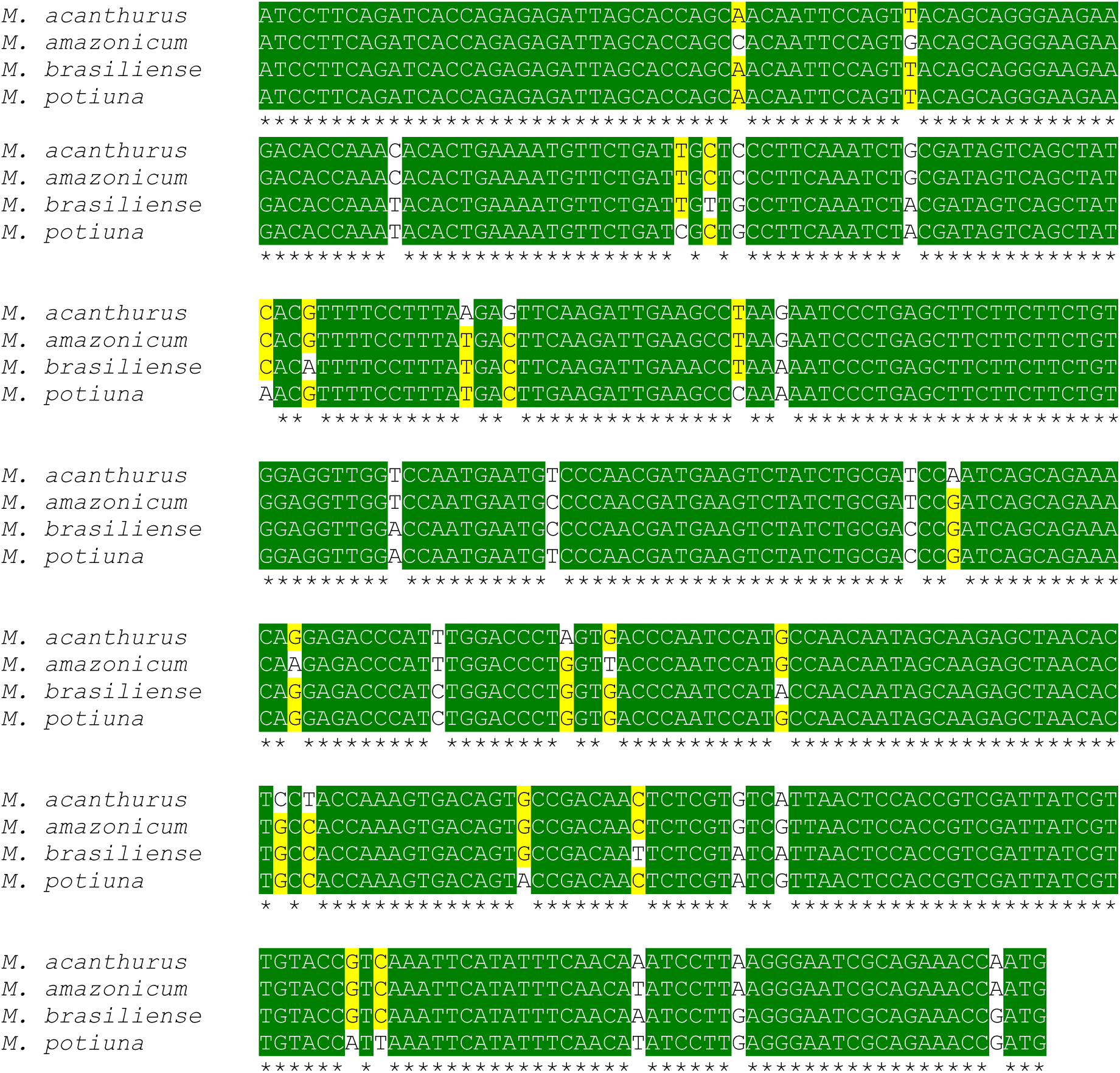
Multiple alignments of partial gene sequences for the gill sodium-potassium-two chloride symporter, amplified by primer pair NKCC_Mk_F1/R1 in several species of freshwater shrimp, *Macrobrachium*. The 415-base pair nucleotide sequences shown are from *Macrobrachium acanthurus* (GenBank deposit MG385140) and *M. amazonicum* (MG385141), both diadromous species, and from *M. brasiliense* (MG385142) and *M. potiuna* (MG385143), both hololimnetic species. Identical bases among all four species are shown in green with three identical bases in yellow.

**Supplementary Figure 2.**
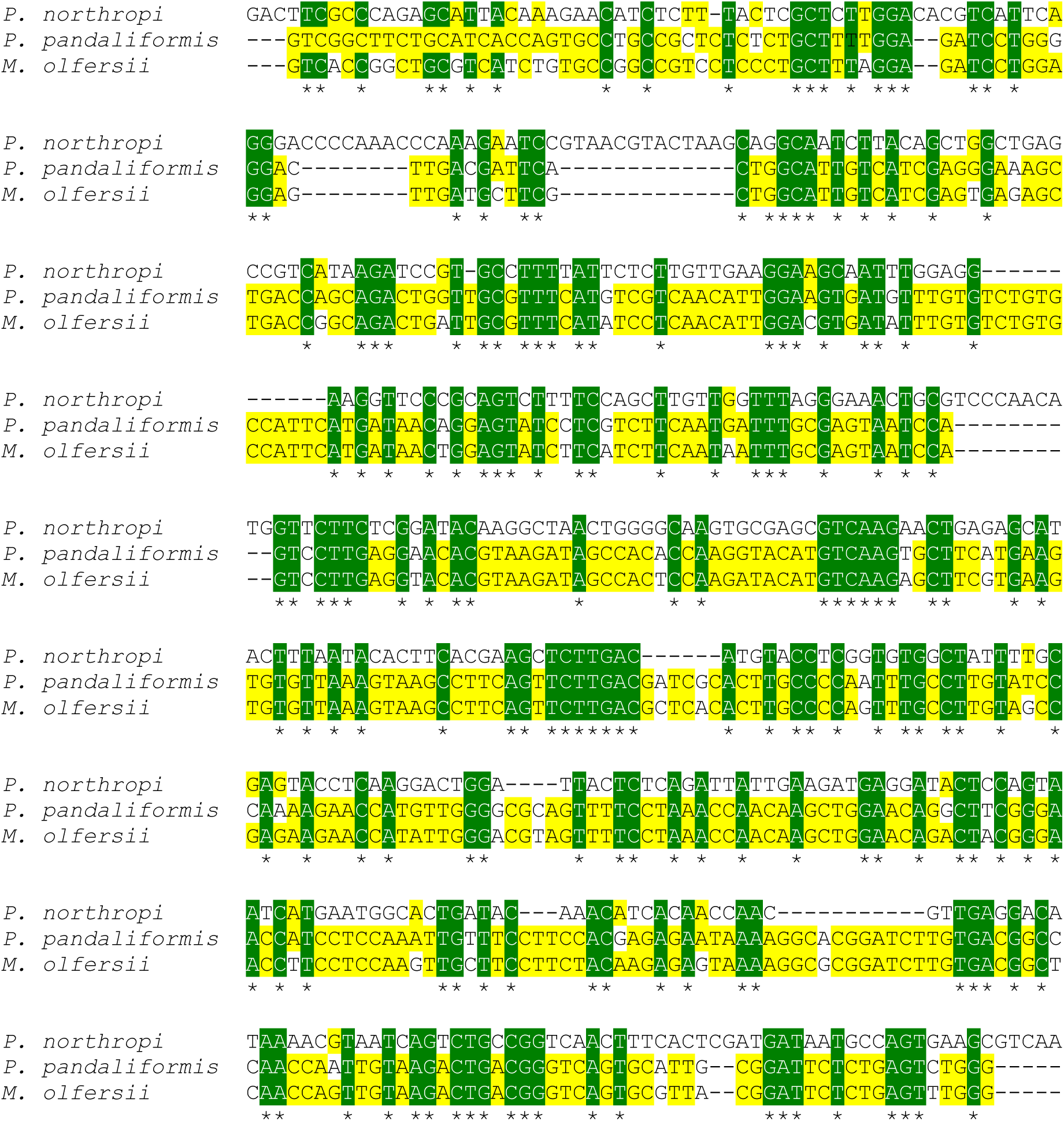
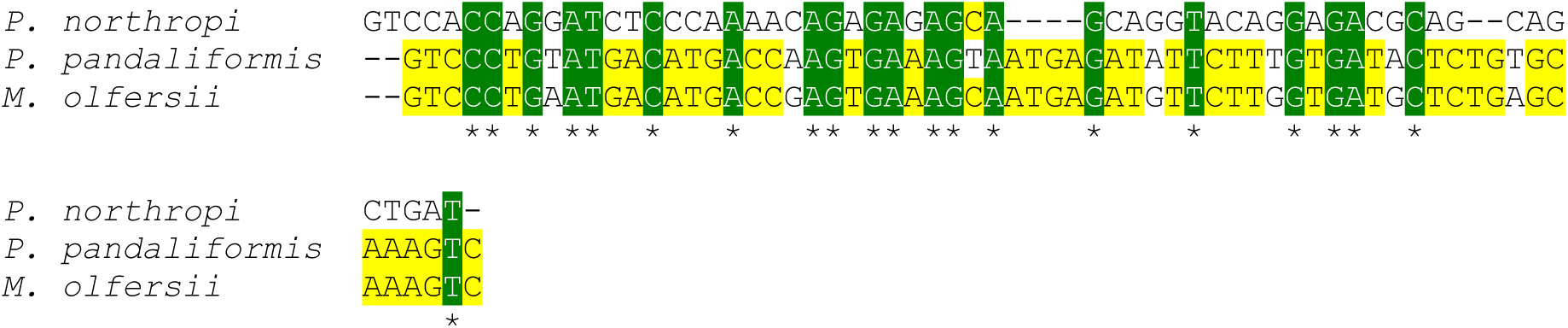
Multiple alignments of partial nucleotide sequences for the gill sodium-potassium-two chloride symporter, amplified by primer pair NKCC_Mk_F2/R2 in three species of palaemonid shrimps. The 561-base pair nucleotide sequences shown are from *Palaemon northropi*, a tide pool shrimp (GenBank deposit MG652470), *P. pandaliformis* (MG652469), an estuarine shrimp, and *Macrobrachium olfersii* (MG652471), a diadromous freshwater shrimp. Identical bases among all three species are shown in green with two identical bases in yellow.

**Supplementary Figure 3.**
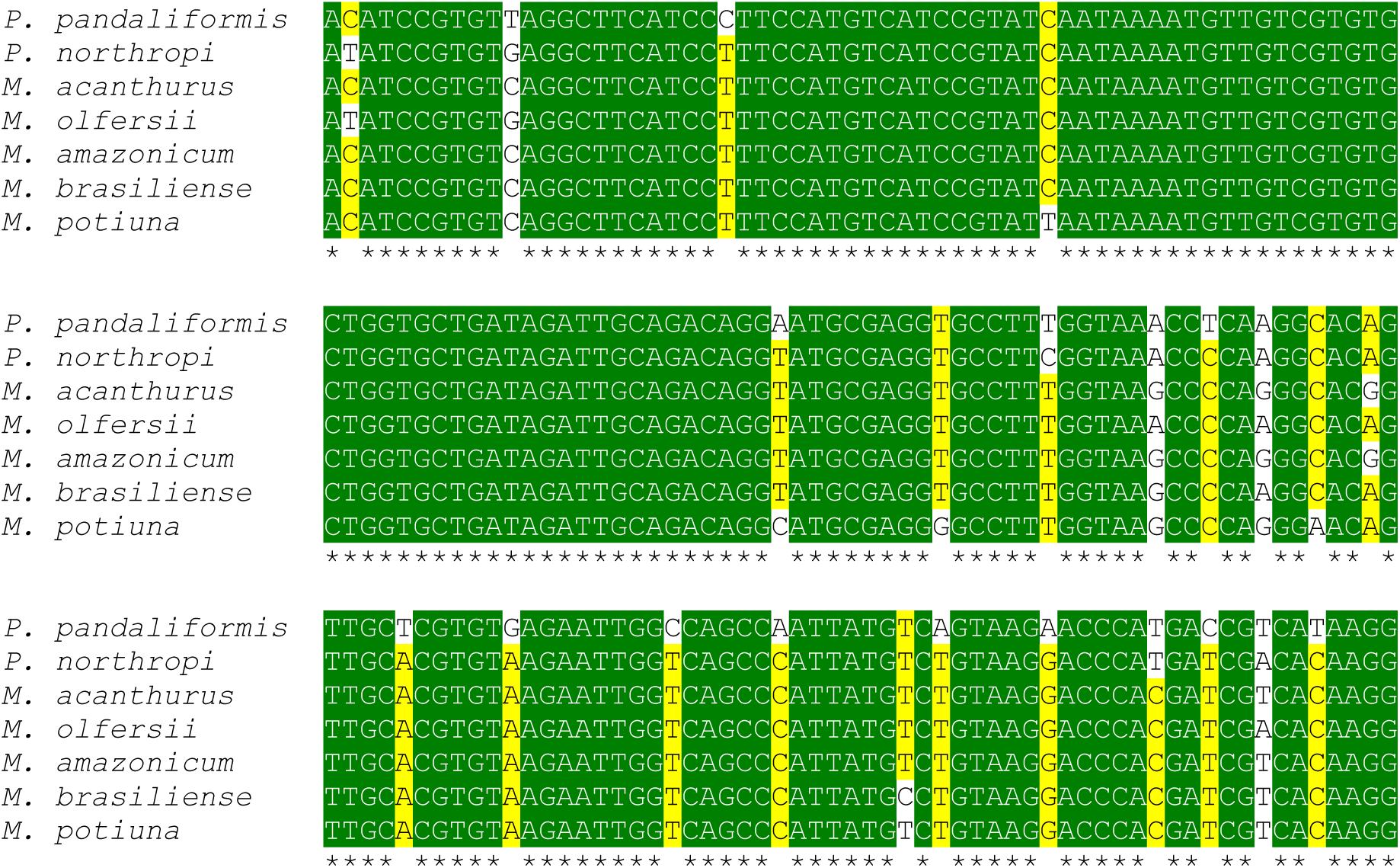
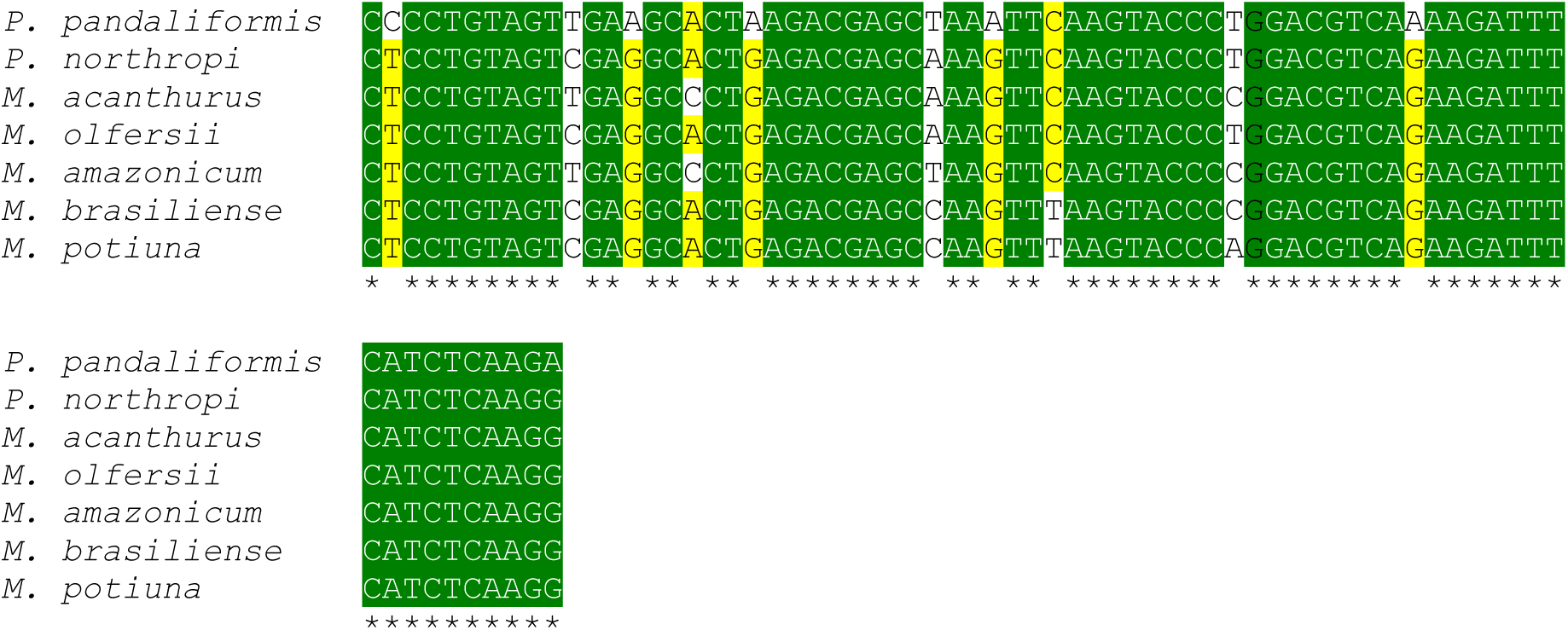
Multiple alignments of partial gene sequences for the gill ribosomal protein L10, amplified by primer pairs RPL10_Cs_F/R and RPL10_Pal_F/R in several species of palaemonid shrimp from different salinity habitats. The 251-base pair nucleotide sequences shown are from *P. northropi*, a tide pool shrimp (Genbank deposit JN25113.5), *P. pandaliformis*, an estuarine shrimp (KP89067.1), *Macrobrachium acanthurus* (JN25113.4), *M. amazonicum* (GU36606.5) and *M. olfersi* (KT78351.5), all diadromous freshwater species, and from *M. brasiliense* (JN25113.3) and *M. potiuna* (KU72624.4), both hololimnetic freshwater species. Identical bases among all seven species are shown in green with five or more identical bases in yellow.

